# Calmodulin requires Ca^2+^ to integrate into the spindle pole body and modulate the cytokinetic ring constriction in fission yeast cells

**DOI:** 10.64898/2026.05.08.723810

**Authors:** Marium Zehra, Debatrayee Sinha, Ajay Sharma, Anusha Gaddam, Jayden Alex Chacko, Qian Chen

## Abstract

Although calmodulin is best known as a calcium sensor, it also possesses Ca^2+^-independent functions, exemplified by the surprising finding that the budding yeast cells remain viable using a calmodulin mutant that can’t bind Ca^2+^ (Geiser et al., 1991). It remains unclear why Ca^2+^-calmodulin is required in other yeasts or vertebrates. Here, we determined whether such holo-calmodulin is an essential cytoskeletal protein in fission yeast. The *S. pombe* calmodulin Cam1 was an integral part of both the spindle pole body (SPB) and many actin structures at the equatorial division plane. Two mutants Cam1-2V and -3V, which bound Ca^2+^ poorly, reduced their localization at the SPBs by 90%. However, their presence in the actin structures remained unchanged. Replacing the endogenous *cam1* with *cam1-2V* cut the number of the Cam1-interacting protein Pcp1 in the SPB by ∼70% and delayed mitosis. In contrast, the assembly and constriction of the cytokinetic ring, which depends on the Cam1-interacting myosins Myo1, Myo51 and Myo52, accelerated. The temperature-sensitive *cam1-2V* mutant was rescued by either over-expression of *pcp1* or deletion of *myo1*. Thus, Cam1 depends on Ca^2+^ to promote the SPB assembly and to modulate the actomyosin ring constriction, suggesting holo-calmodulin as an essential cytoskeletal protein.

## INTRODUCTION

Calmodulin is one of the most evolutionarily conserved proteins in eukaryotes. First discovered as a Ca^2+^-dependent activator of phosphodiesterase (Cheung, 1970; Lin et al., 1974), this Ca^2+^-binding protein has since been found in almost all plants, fungi, and animals (Halling et al., 2016; Mohanta et al., 2017). All calmodulins share a highly conserved sequence of less than 200 amino acids. It consists of two pairs of EF-hands, each of which has a signature helix-loop-helix secondary structure. Each EF-hand can bind one Ca^2+^ through electrostatic interactions. Binding Ca^2+^ transforms calmodulin from apo- to holo-form, capable of binding many of its targets including calmodulin-dependent kinases (Babu et al., 1985). Mutations of the human calmodulin, of which there are three copies in the genome (Halling et al., 2016), are linked to many genetic disorders including ventricular tachycardia and sudden cardiac death (Crotti et al., 2013, 2019; Nyegaard et al., 2012).

Despite the common perception of calmodulin solely as an intracellular calcium sensor, its essential functions are completely Ca^2+^-independent in the unicellular model organism *S. cerevisiae*. This budding yeast possesses only one essential calmodulin Cmd1 that has been extensively studied (Davis et al., 1986). Surprisingly, the yeast is viable without significant growth defects even with a calmodulin mutant that can’t bind Ca^2+^(Geiser et al., 1991). Instead, the interactions between Cmd1 with its essential targets of the cytoskeletal proteins are all Ca^2+^-independent. They include both type I and V myosins (S. Brockerhoff et al., 1994; Geli, 1998) and the pericentrin-like protein Spc110 in the yeast microtubule-organizing center of the spindle pole body (SPB) (Geiser et al., 1993; Stirling et al., 1994). The studies of the budding yeast Cmd1 suggest that calmodulin can carry out its essential functions independent of Ca^2+^ in unicellular eukaryotic organisms.

The fission yeast *S. pombe* calmodulin Cam1 shares some similarities with its *S. cerevisiae* orthologue, but it has diverged substantially. As in budding yeast, fission yeast Cam1 is an essential protein (Moser et al., 1997), serving as a constitutive component of both the microtubule-nucleating SPB and actin-based cytoskeletal structures, such as endocytic patches and the cytokinetic contractile ring (Eng et al., 1998; Moser et al., 1997a). Cam1 interacts with the Spc110 homologue Pcp1 (Bestul et al., 2017a; Chen et al., 2024; Flory et al., 2002; Rosenberg et al., 2006) and both type I and V myosins (Lee et al., 2000; Win et al., 2001).

However, there are also substantial differences between calmodulins from these two yeasts that are separated by more than 300 million years of evolution (Rhind et al., 2011). To start, the fission yeast calmodulin binds four Ca^2+^ ions, like the vertebrate ones, but the budding yeast calmodulin binds only three (Moser et al., 1995). Not surprisingly, the fission yeast protein exhibits a much higher sequence identity (75%) with the vertebrate calmodulin than with that of budding yeast (57%) (Takeda & Yamamoto, 1987). Likely as a result of their substantial differences, the fission yeast calmodulin can’t be replaced by the budding yeast Cmd1, but it can be replaced with the vertebrate calmodulin (Moser et al., 1995). Most importantly, in contrast to the budding yeast calmodulin, the Ca^2+^-binding capacity of fission yeast Cam1 is indispensable for cell viability (Moser et al., 1995), but the reason remains unclear.

Here, to address this long-standing question of why calmodulin needs Ca^2+^ for its essential functions in fission yeast, we determined whether the distribution of Cam1 between actin- and microtubule-based cytoskeletal structures depends on Ca^2+^. We made two Cam1 mutants, each with a substantial defect in binding Ca^2+^. Both mutants, Cam1-2V and -3V, were largely absent from the SPBs, despite their normal localization at the actin-based structures. When the endogenous gene was replaced by *cam1-2V*, the number of Pcp1 molecules at the SPBs dropped by 70% while the other SPB proteins changed little. This led to strong temperature-sensitive defects in mitosis including loss of SPBs and a delay in mitosis. In contrast, the localization of both type I and V myosins, Myo1, Myo51 and Myo52 in the actin cytoskeletal structures, remained unchanged in this *cam1* mutant. Unlike the delayed mitosis, the *cam1-2V* mutation accelerated cytokinesis, particularly the contractile ring assembly and constriction. The temperature-sensitive growth of *cam1-2V* mutant was rescued by either over-expression of *pcp1* or deletion of *myo1*. Overall, our results show that fission yeast Ca^2+^-Cam1 is an essential component of both actin and microtubule cytoskeletal structures in cell division, suggesting that Ca^2+^ bound calmodulin is required for its paradoxical effect towards the microtubule and actin cytoskeletal structures in fission yeast and vertebrates.

## RESULTS

### Ca^2+^-binding mutant of Cam1 fail to integrate into the SPB

To determine whether the intracellular distribution of Cam1 depends on Ca^2+^, we first determined the localization of Cam1 through fluorescence microscopy. When tagged with GFP at its N-terminus, the endogenously expressed GFP-Cam1 was localized to the SPBs throughout the cell cycle. In addition, it was found at the actin patches at the cell tips, as well as the equatorial division plane (Fig. 1A). This is consistent with the previous reports (Moser et al., 1997). In addition, our 3D reconstruction of the division plane showed GFP-Cam1 as numerous puncta in a donut-shaped disk surrounding the actomyosin contractile ring (Supplemental Fig. S1A). This disk expanded gradually towards the cell center during cytokinesis as the ring constricted. We concluded that the endogenous Cam1 is a component of both microtubule-organizing center of SPB and the actin-based structures at both the cell tip and division plane.

**Figure 1.**
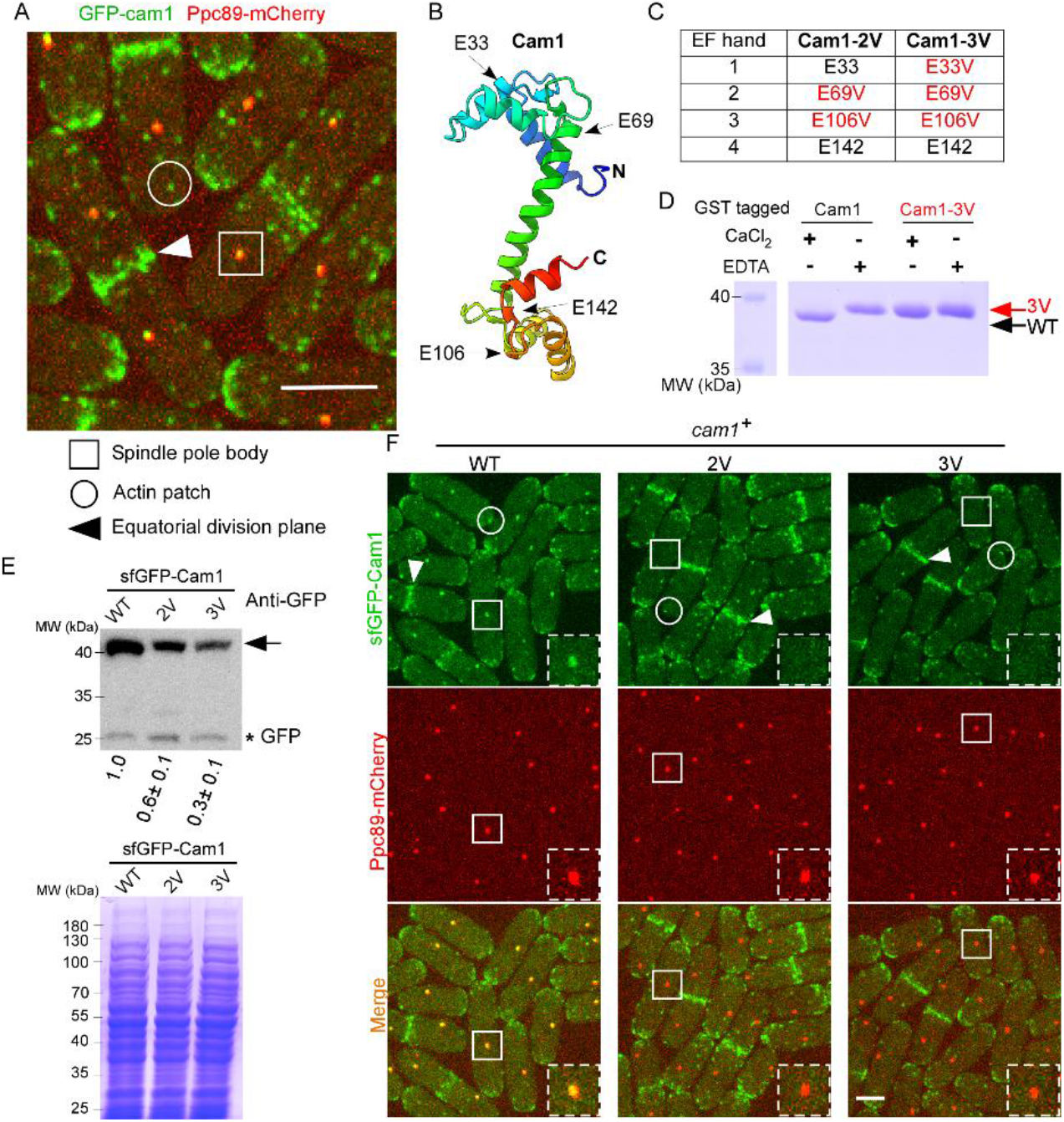
Ca^2+^-binding mutants of fission yeast calmodulin Cam1-2V and -3V are mostly absent from SPBs. (A) Micrograph of the cells co-expressing GFP tagged endogenous Cam1 (green) and the SPB marker Ppc89-mCherry (red). GFP-Cam1 was localized to the SPBs (box), actin patches (circle) at the cell tip and the division plane (arrowhead). (B) Ribbon diagram (rainbow-colored) of the AlphaFold3 predicted tertiary structure of Cam1. Arrow: The Ca^2+^-interacting glutamine acids (E) in each of the four EF-hands. (C) A list of the glutamine acid (E) to valine (V) mutations in the mutants Cam1-2V and -3V. (D) Scanned Coomassie Blue-stained SDS-PAGE gel for electrophoresis of the recombinant GST tagged Cam1 and Cam1-3V proteins in the presence of either 4mM CaCl_2_ or 2mM EDTA. (E) Scanned anti-GFP blot (top) and the corresponding Coomassie Blue-stained SDS-PAGE gel (bottom) of the lysates of the yeast cells expressing sfGFP-Cam1 (WT), sfGFP-Cam1-2V (2V), and sfGFP-Cam1-3V (3V) respectively. Arrow: sfGFP-Cam1. Asterisk: GFP. Number: Normalized intensity value (average±standard deviations, N=3) of the mutant Cam1 relative to the wild-type one. Left: molecular weight standard. (F) Micrographs of the *wild-type* cells co-expressing Ppc89-mCherry (red) and sfGFP tagged the *wild-type* Cam1, 2V, and 3V mutants respectively (green). Dashed Box: magnified view of representative SPB (Box). Representative data from three biological repeats is shown. Bar: 5µm.

To block Cam1 from binding to Ca^2+^, we targeted its four EF-hand motifs for mutagenesis (Fig. 1B), based on a strategy first used by Trisha Davis’s group (Moser et al., 1995). Each EF-hand loop of Cam1, consisting of 12 amino acids, ends with a highly conserved glutamic acid (E) (Fig. 1B). Replacing this negatively charged residue with a non-charged one, valine (V), inhibits binding of Ca^2+^ (Geiser et al., 1991). We constructed two mutants, *cam1-E69V E106V (2V)* and *cam1-E33V E69V E106V (3V),* by replacing either two or three of the four glutamic acids (Fig. 1C). We tested the *in vitro* binding of Cam1-3V to Ca^2+^, which is expected to be weaker than that between Cam1-2V and Ca^2+^. We purified the recombinant wild-type Cam1 and Cam1-3V from bacteria (Fig. 1D). Consistent with the previous studies (Davis et al., 1986), the recombinant Cam1, even tagged with GST, migrated significantly more quickly in the presence of Ca^2+^ during electrophoresis than it did in the absence of Ca^2+^ (Fig. 1D). In contrast, the mutant Cam1-3V migrated similarly either with or without Ca^2+^, confirming its expected deficiency in binding to Ca^2+^ (Fig. 1D). We concluded that both 2V and 3V mutants of Cam1 bind Ca^2+^ with lower affinity than that of the wild-type protein.

To determine the intracellular distribution of these two Cam1 mutants, we tagged their N-terminus with super-fold GFP (sfGFP). We first tried to replace the endogenous *cam1* gene with these sfGFP tagged mutants. However, this failed to produce viable yeasts, likely due to the lethality of these alleles. As an alternative strategy, we expressed them and the wild-type sfGFP-Cam1 at the exogenous *ura4* locus driven by the constitutive actin promoter Pact1 is of similar strength as the endogenous *cam1* promoter (Supplemental Fig. S1B). The exogenously expressed *cam1* mutants did not alter the growth of the yeasts (Supplemental Fig. S1C). However, the mutant proteins, both 2V and 3V, were ∼40% and 70% less abundant respectively than the wild-type sfGFP-Cam1, based upon the anti-GFP immunoblots (Fig. 1E). As expected, the exogenously expressed sfGFP-Cam1 was found at the SPBs, the cell tips and the division plane (Fig. 1F). In contrast, the mutants, both 2V and 3V, were largely absent from the SPBs (Fig. 1F and Supplemental Fig. S1D), even though both remained present at the actin structures at the cell tips and the division plane (Fig. 1F). We concluded that Ca^2+^-binding mutations of Cam1 significantly reduce their localization at SPBs and their *in vivo* stability.

Next, we decided to precisely measure the amount of these Cam1 mutants in the SPBs throughout cell division using quantitative fluorescence microscopy (Fig. 2A). We used the movement of SPBs, marked by Ppc89-mCherry, as a “clock” for mitosis (Wu et al., 2003). The separation of SPBs was designated as time zero. During mitosis, the fluorescence intensity of the wild-type sfGFP-Cam1 at the SPBs gradually increased by about two folds (+93%, P<0.001). It started 10 mins before the SPB separation (-10 min) before plateauing at +10 min (Fig. 2B). In comparison, the fluorescence intensities of either 2V or 3V mutants at the SPBs were already ∼90% lower than that of the wild-type protein before the mitotic entry (Fig. 2A and B). Furthermore, the SPB localization of the mutants failed to increase significantly during this period of between -10 and +10 min (P<0.001), as the wild-type protein did. Therefore, the two mutants reduce the mitotic SPB localization of Cam1 by 90% respectively.

**Figure 2.**
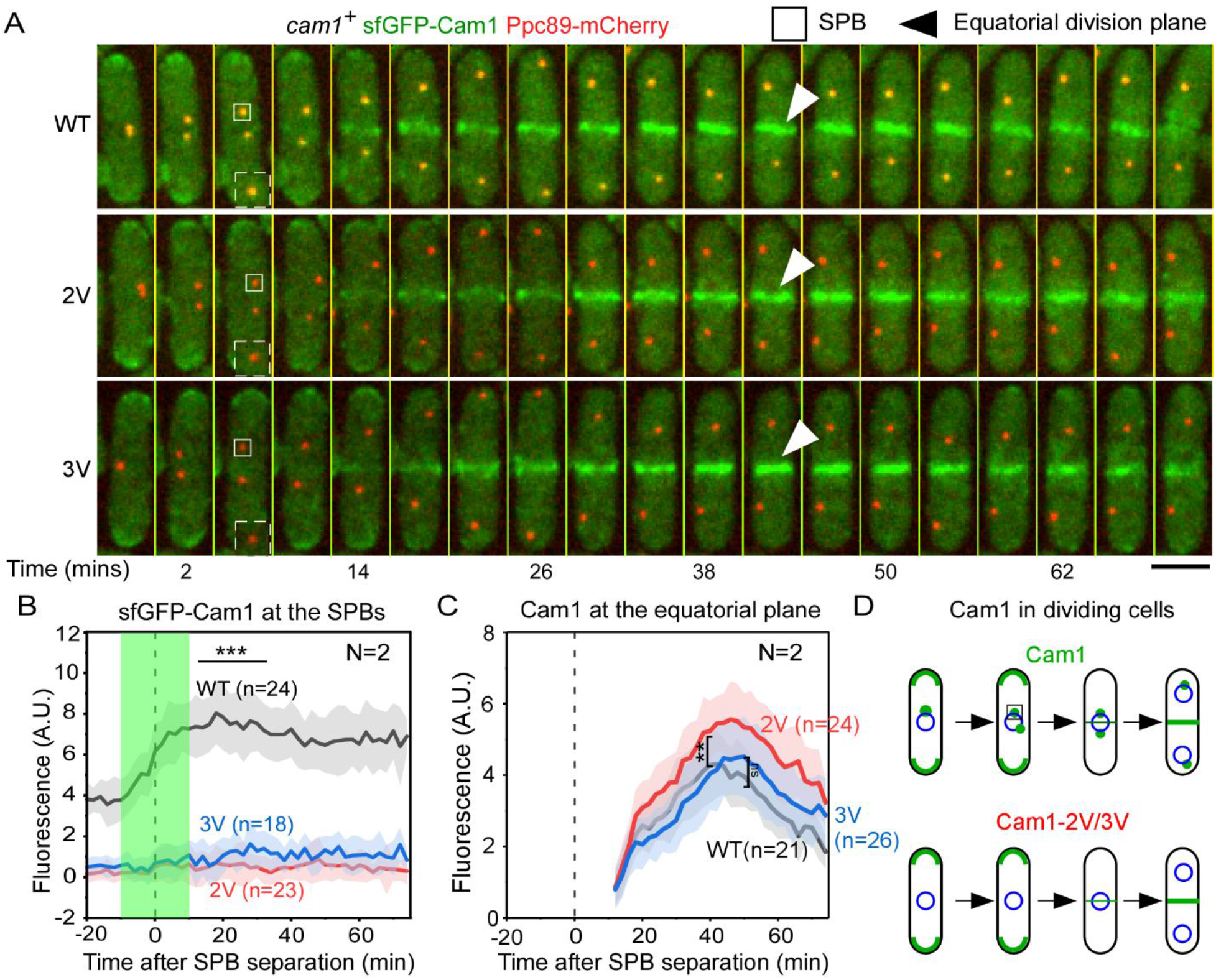
Both cam1-2V and -3V mutations block 90% of Cam1 molecules from the mitotic SPBs, but they have no effect on the localization of Cam1 at the equatorial division plane. (A) Time series of dividing cells co-expressing Ppc89-mCherry (red) and either sfGFP-Cam1 (green, WT), or -2V, or -3V. Dashed Box: magnified view of representative SPB (Box). Arrowhead: the equatorial division plane. Number: time in minutes after the separation of SPBs. (B-C) Average time courses of the fluorescence intensities of sfGFP-Cam1 (WT, gray), -2V (red), and -3V (blue) at either the SPBs (B) or the equatorial division plane (C). Dashed line: Time of the SPB separation. Shade: the period during which the amount of GFP-Cam1 at the SPB doubled. (D) A diagram summarizing the intracellular localization of Cam1 and its mutants throughout the cell cycle. Representative data from two independent biological repeats is shown. ***: P<0.001, **: P<0.01, N.S: P>0.05, based on two-tailed student’s t-tests. Bar: 5µm.

We considered two potential explanations for this result. The reduced presence of the Cam1 mutants at the SPBs could be attributed to either their inability to bind to Ca^2+^ or their reduced protein stability. If the latter was the primary cause, we expected that the mutants’ localization to the equatorial plane of the dividing cells would decline by a similar extent as they did in the SPBs. To test this, we measured the localization of the wild-type Cam1 and the two mutants at the equatorial plane throughout cell division. The wild-type sfGFP-Cam1 first appeared at the division plane at +12 min (Fig. 2A). Its fluorescence intensity gradually increased ∼5-fold over the next 30 mins, peaking at +40 min (Fig. 2C). In comparison, the two mutants appeared at the division plane at the same time. Their fluorescence intensities at the division plane peaked at +46 and +50 min, respectively. The peak fluorescence intensities of the wild-type and even the 3V mutant at the division plane were similar to each other (P>0.05), both just slightly lower than that of the 2V mutant (P<0.01). This result demonstrated that Cam1 mutants remain a normal part of the actin structures at the equatorial plane despite their reduced stability. Their reduced capability to bind Ca^2+^ more likely explains these mutant proteins. Therefore, binding to Ca^2+^ is required for Cam1 to integrate into the SPBs, but it is dispensable for Cam1 to be a part of the actin cytoskeletal structures at the equatorial division plane (Fig. 2D).

### The SPB localization of Pcp1 is reduced significantly in the mutant *cam1::cam1-2V*

We determined the cellular effects of displacing Cam1 from the SPB. We replaced the endogenous gene with either *cam1-2V* or *-3V* without GFP. We successfully created *cam1::cam1-2V*, but not *cam1::cam1-3V,* despite repeated efforts, suggesting that the latter may be a lethal allele. Our 2V mutant carried identical point mutations in the *cam1* gene as the existing mutant *cam1-E14* (Moser et al., 1995), except that our mutant allele was linked to two auxotrophic markers, *his3^+^* and *ura4^+^*, as a result of the two-step gene replacement process (Tang et al., 2011). We used our own *cam1::cam1-2V* for the rest of this study, which we henceforth refer to as *cam1-2V* to avoid confusion. Like *cam1-E14*, *cam1-2V* mutant was temperature-sensitive, inviable at 36°C (Fig. 3A).

**Figure 3.**
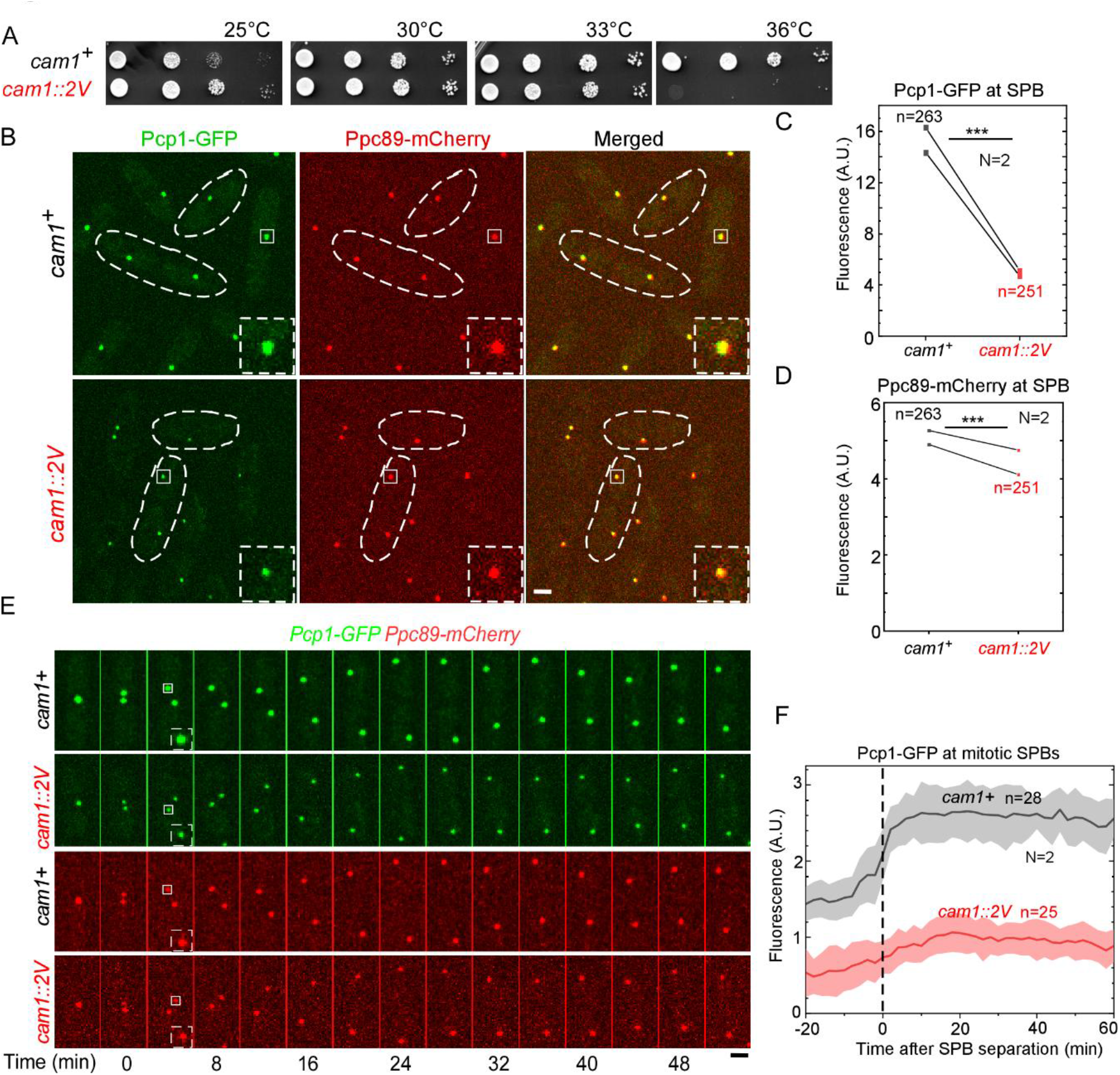
Replacing the endogenous *cam1* with *cam1-2V* reduces the number of pericentrin- like Pcp1 molecules at the SPB by 70%. (A) Ten-fold dilution series of yeasts. The *cam1::cam1-2V* mutant was inviable at 36°C. (B) Micrographs of the *wild-type* and the *cam1::cam1-2V* cells co-expressing both Pcp1-GFP (green) and the SPB marker Ppc89-mCherry (red) at the permissive temperature. Dashed box: magnified view of representative SPB (Box). (C-D) Paired dot plots of the average fluorescence intensities of Pcp1-GFP (C) and Ppc89-mCherry (D) at the SPBs of the *wild-type* and *cam1-2V* mutant cells. (E) Time series of a dividing *wild-type* and *cam1-2V* cell, respectively, expressing both Pcp1-GFP and Ppc89-mCherry. Number: time in minutes after the SPB separation. (D) Average time courses of the fluorescent intensities of Pcp1-GFP at the SPBs of *wild-type* and *cam1-2V* mutant cells, respectively. Cloud: standard deviations. Data is pooled from two independent biological repeats. ***: P<0.001, based on two-tailed student’s t-tests. Bar: 5µm.

We first carried out fluorescence microscopy to characterize the molecular composition of the SPBs of *cam1-2V* at the permissive temperature. We expected that the 2V mutation would have an effect on the SPBs even in this viable condition. First, we measured the localization of the GFP tagged Pcp1 (Fig. 3B), which interacts with Cam1 directly (Chen et al., 2024; Moser et al., 1997a). In an asynchronous population of the *cam1-2V* cells, the average fluorescence intensity of Pcp1-GFP at the SPBs, marked by Ppc89-mCherry, was 68% lower than that of the wild-type protein (P <0.001) (Fig. 3C). In comparison, the fluorescence intensity of Ppc89-mCherry at the SPBs of the mutant cells decreased far less by merely 13% (P <0.001) (Fig. 3D). Next, we quantified the amount of Pcp1-GFP at the SPBs throughout mitosis using time-lapse microscopy, again by using the SPB separation as “time zero” to align the time courses (Fig. 3E). As expected, the fluorescence intensities of Pcp1-GFP at the mother SPBs of the mutant cells, prior to mitosis, was already 68% lower than that of the wild-type cells (Fig. 3E and F). Surprisingly, in the *cam1-2V* cells, the amount of Pcp1-GFP at the SPB did increase gradually by two folds from -10 min till plateauing +10 min, similar to the wild-type cells (Fig. 3F). Toward the end of mitosis, the combined fluorescence intensity of the Pcp1-GFP in two the daughter SPBs of the mutant cells remained 70% lower than that of the *wild-type* (Fig. 3F). We concluded that *cam1-2V* mutation blocks a majority of Pcp1 molecules from integrating into the SPB.

Since *cam1-2V* is temperature-sensitive, we next determined how the molecular composition of the SPBs is altered by the 2V mutation at the restrictive temperature. In addition to Pcp1 and Ppc89, we measured the amount of three other proteins, Cdc11, Alp4, and Plo1, in the SPBs of the asynchronized cells at 36°C using quantitative fluorescence microscopy. Cdc11 is the fission yeast centriolin ortholog (Tomlin et al., 2002). Alp4 is one of the six components of the fission yeast γ-tubulin complex that nucleate mitotic spindles (Masuda et al., 2013). Plo1 is an essential mitotic kinase recruited to the SPBs by Pcp1 (Fong et al., 2010). As expected, the fluorescence intensity of Pcp1-GFP at the SPBs of the *cam1* mutant cells decreased by 71% at the restrictive temperature, compared to the *wild-type* cells (Fig. 4A). In comparison, the fluorescence intensity of Cdc11-GFP at SPBs did not change in the *cam1-2V* mutant cells (Fig. 4A). Instead of a decline, the *cam1* mutation increased the SPB localization of Ppc89-mCherry slightly (15%) (Fig. 4A). Lastly, both Alp4-GFP and Plo1-GFP decreased slightly in the SPBs (6 and 20% respectively) of *cam1-2V* cells (Fig. 4A and Supplemental Fig. S2A-B). In addition to the altered molecular composition of SPBs, we also found that an average of 17% of the mutant cells possessed no detectable SPBs, that were marked by either Ppc89-mCherry or Cdc11-GFP (Fig. 4B-E and Supplemental Fig. S2C). Not surprisingly, these SPB-less cells failed to assemble mitotic spindles (Supplemental Fig. S2C). Next, we determined whether Ca^2+^-Cam1 has a direct role in the dynamics of the mitotic spindles through time-lapse microscopy. Many mutant cells failed to progress to mitosis at the restrictive temperature. For the few that did enter mitosis, most of them (90%) assembled, extended, and disassembled their mitotic spindle normally, similar to the *wild-type* cells (P>0.05) (Supplemental Fig. S3A-C). Only a small fraction (10%) of the mutant cells assembled either bipolar or multi-polar spindles that failed to extend fully (Supplemental Fig. S3A and C). We concluded that Ca^2+^-Cam1 is essential for the SPB assembly, but it only has an indirect role in regulating turnover of the mitotic spindles.

**Figure 4.**
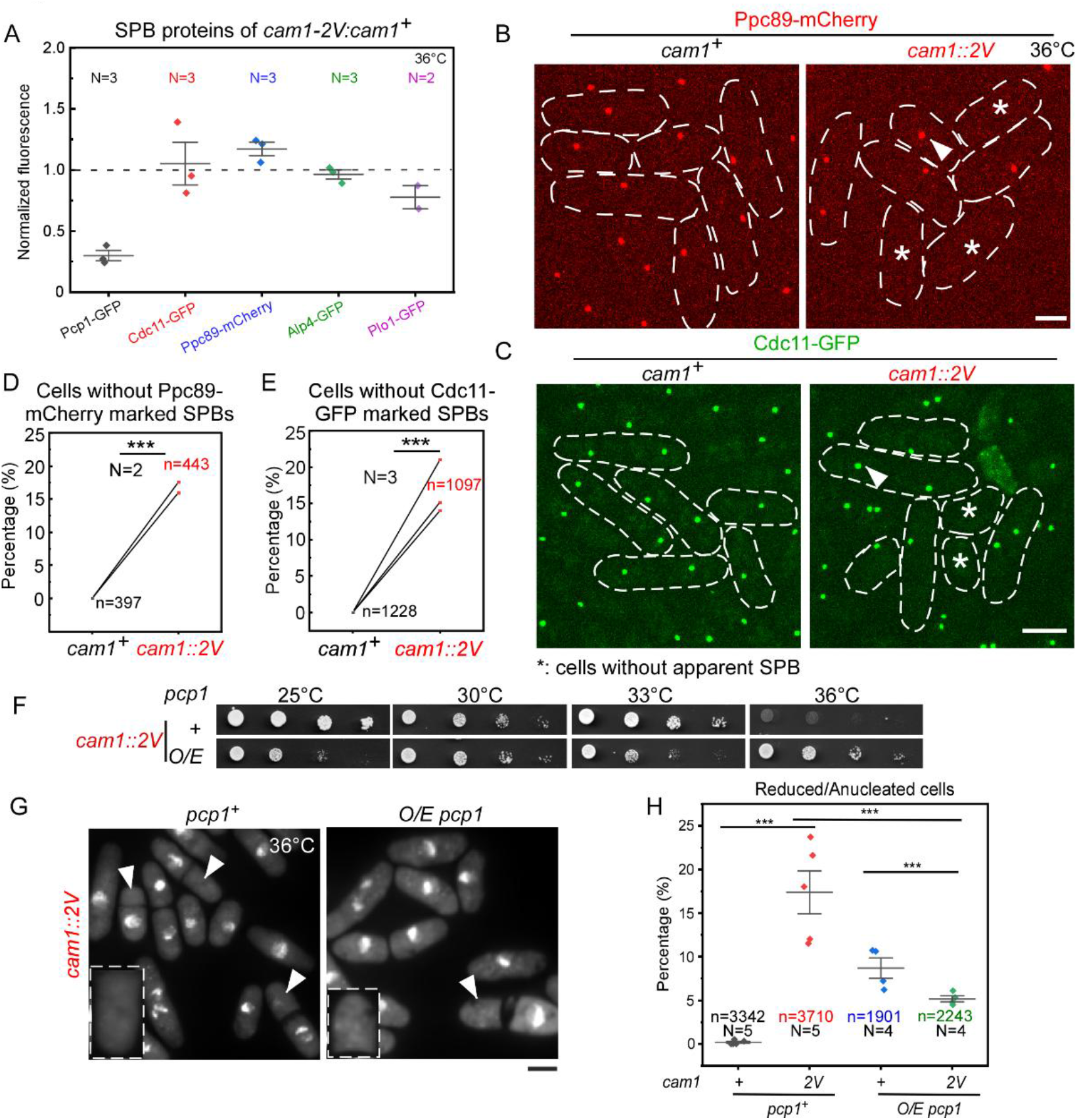
The loss of Pcp1 leads to SPB assembly defects in *cam1-2V* mutant at the restrictive temperature. (A) Dot plot of average fluorescence intensities of five proteins at the SPB of the *cam1-2V* mutant at 36°C. The values were normalized against that of the *wild-type*. Data is pooled from three biological repeats except Plo1-GFP. (B-C) Micrographs of the *wild-type* and *cam1-2V* cells expressing either Ppc89-mCherry (red, B) or Cdc11-GFP (green, C) at 36°C, respectively. Arrowhead: SPBs. Asterisk: representative mutant cells without a detectable SPB marked by either Ppc89-mCherry or Cdc11-GFP. (D-E) Dot plots of the percentage of the mutant cells without a SPB, based on the fluorescence of either Ppc89-mCherry (D) or Cdc11-GFP (E). (F) Ten-fold dilution series of yeasts. Overexpression (O/E) of *pcp1* rescued the temperature-sensitive growth of *cam1-2V* at 36°C. (G) Micrographs of DAPI-stained cells at 36°C. Arrowhead: representative cells without a visible nucleus. Dashed Box: magnified view of representative cells. (H) Dot plots of the fraction of anucleate cells. Brackets: average ± standard deviation. ***: P<0.001, based on two-tailed student’s t-tests.

To determine whether loss of Pcp1 from the SPBs alone underlies the temperature-sensitive lethality of *cam1-2V*, we tested whether a hypermorphic *pcp1* mutant could rescue this *cam1* mutant. We over-expressed Pcp1 by replacing its endogenous promoter with the strong actin promoter Pact1 and an N-terminal GFP. Consistent with a previous report (Flory et al., 2002), over-expressed GFP-Pcp1 formed several large intracellular puncta in addition to its expected localization to the SPBs (Supplemental Fig. S4A). This hypermorphic *pcp1* mutation largely rescued the growth of *cam1-2V* mutant at 36°C (Fig. 4F). It reduced the fraction of *cam1-2V* cells without an apparent nucleus from 17% to 5% (P <0.001) (Fig. 4G-H). Thus, the loss of Pcp1 from the SPBs is a primary cause of the lethality of *cam1-2V* mutation at the restrictive temperature.

### Ca^2+^-Cam1 modulates the actomyosin contractile ring constriction and promotes the daughter cell integrity during cytokinesis

We next determined whether the actin cytoskeletal structures are altered by the *cam1-2V* mutation. First, we examined the localization of the Cam1-interacting unconventional myosins, Myo51, Myo52 and Myo1, in *cam1-2V* mutant cells at either permissive or restrictive temperatures. First, Myo52, tagged with tdTomato at the endogenous locus, was localized normally to the cell tips and the equatorial division plane of *cam1-2V* mutant cells at either temperature (Fig. 5A and Supplemental Fig. S5A). This was similar to the *wild-type* cells (Fig. 5A and Fig S5A). Secondly, the other type V myosin Myo51, similarly tagged with tdTomato at its endogenous locus, remained a part of the contractile ring in the *cam1* mutant cells, unchanged from the *wild-type* cells (Fig. 5A and Supplemental Fig. S5A). Lastly, the type I myosin Myo1, tagged with GFP endogenously, was found at numerous endocytic actin patches in *cam1-2V* cells like the *wild-type* cells (Fig. 5A and Supplemental Fig. S5A). Therefore, we concluded that none of the unconventional myosins require Cam1-Ca^2+^ for their intracellular localization, unlike Pcp1.

**Figure 5.**
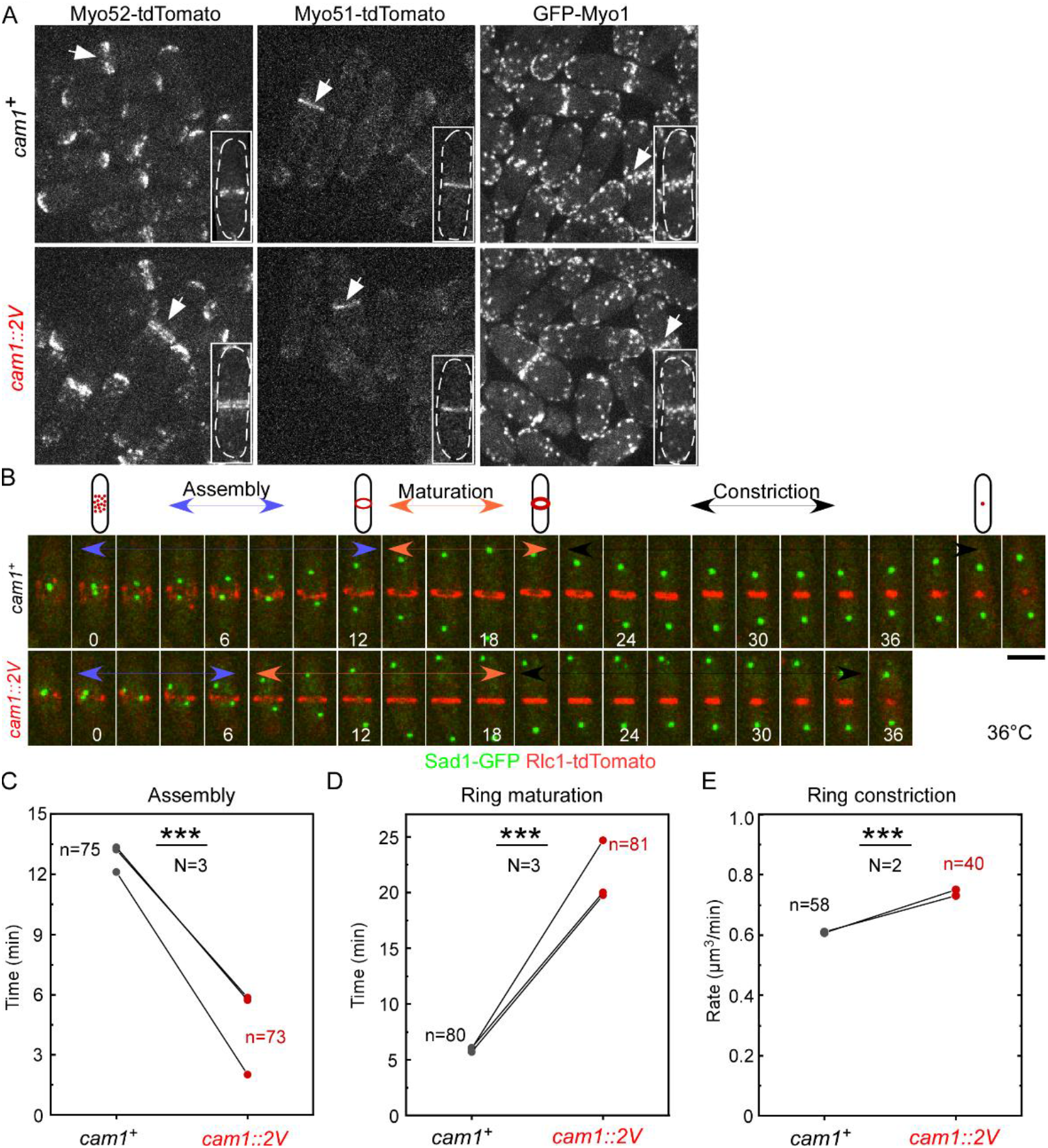
cam1-2V does not change the localization of unconventional myosins, but it accelerates the cytokinetic ring assembly and constriction at the restrictive temperature. (A) Micrographs of the *wild-type* (top) and *cam1-2V* (bottom) cells expressing either Myo52-tdTomato, Myo51-tdTomato, or GFP-Myo1 at 36°C. Arrow: the equatorial division plane. Box: magnified view of representative cells. (B) Top: cartoon diagram summarizing the steps of fission yeast cytokinesis. Bottom: time-series of a dividing *wild-type* (top) and *cam1-2V* (bottom) cell co-expressing the SPB marker Sad1-GFP (green) and the contractile ring marker Rlc1-tdTomato (red) at 36°C. Number: time in minutes after the SPB separation. (C-E) Paired dot plots of the durations of the contractile ring assembly (C), maturation (D), and the rate of the ring constriction (E) in the *wild-type* and *cam1-2V* cells. Data is pooled from three independent biological repeats. ***: P<0.001, based on two-tailed student’s t-tests.

Next, we determined whether Ca^2+^-Cam1 regulates the actin cytoskeletal structures at the division plane. Both type V myosins, Myo51 and Myo52, promote the assembly and constriction of the contractile ring (Laplante et al., 2015). Therefore, we measured the duration of assembly and maturation of the actomyosin contractile ring marked by the myosin regulatory light chain Rlc1-tdTomato, as well as the constriction rate of the ring (Fig. 5B). At the permissive temperature, *cam1-2V* mutant cells assembled the ring more quickly than the *wild-type* cells did (Supplemental Fig. S5B), but their rings took longer to mature (Supplemental Fig. S5C). The mutant cells constricted their rings at the same rate as the *wild-type* cells did (Supplemental Fig. S5D). At the restrictive temperature, the *cam1-2V* cells were much quicker (4±14 vs. 13±2 mins, P<0.001) to assemble the contractile ring than *wild-type* cells did (Fig. 5B-C). Following the assembly of the contractile ring, the *cam1* mutant cells took longer than the *wild-type* cells (∼21±14 vs. 6±1 mins, P<0.001) to initiate the contractile ring constriction (Fig. 5B and D). However, the contractile ring in *cam1-2V* mutant cells constricted at a 20% higher rate than that of the *wild-type* (0.74±0.12 vs. 0.61±0.05 µm/min, n=40, P<0.001) (Fig. 5E). Lastly, we determined whether Ca^2+^-Cam1 contributes to the separation of daughter cells, the last step of cytokinesis following the closure of the contractile ring. At the restrictive temperature, the *cam1-2V* mutant exhibited a similar septation index to that of *wild-type* cells (Supplemental Fig. S6A-B). Few (<1%) of the *cam1* mutant cells were multi-septated (0.6±0.5%). Only a small fraction (4.5±1.5%) of the *cam1-2V* mutant cells deposited septum ectopically at either the cell tip or the side (Supplemental Fig. S6A and C). Fluorescence microscopy revealed that the daughter cells of *cam1-2V* mutant took a similar amount of time to separate as the *wild-type* cells did (11±5 vs. 12±2 mins, n=37 and 58, N=2, P>0.05) (Supplemental Fig. S6D-E). However, unlike the *wild-type* daughter cells, a significant portion (11±1%, n=235) of the *cam1-2V* daughter cells lysed within two minutes of separation (Supplemental Fig. S6F). Overall, we concluded that Ca^2+^-Cam1 modulates both the contractile ring assembly and constriction, and it is also critical for the integrity of daughter cells during cytokinesis.

Lastly, we tested whether there is any interplay between the mutants of *cam1* and the three unconventional myosins. We tested the genetic interactions between *cam1-2V* and *myo1Δ, myo51Δ,* and *myo52Δ*, respectively. Neither *myo52Δ* nor *myo51Δ* exhibited a genetic interaction with *cam1-2V*. This was in stark contrast to the strong negative genetic interaction between *cam1-2V* and *pcp1-15* (Fig. 6A). In comparison, *myo1Δ* exhibited a strong positive genetic interaction with the *cam1* mutant. Deletion of *myo1* rescued the temperature-sensitivity of *cam1-2V* at 36°C (Fig. 6A). We hypothesized that this may be due to an influx of Cam1 to the SPBs when Cam1 is freed from the actin patches in the absence of Myo1. The fluorescence intensities of GFP-Cam1 at the SPBs of the *myo1* mutant cells increased by almost three folds (Fig. 6B-C) from that of the *wild-type* cells. In contrast, the fluorescence intensities of Ppc89-mCherry at the SPB only decreased slightly in the mutant cells (Fig. 6C). Therefore, Myo1 modulates the SPB localization of Ca^2+^-Cam1, providing a potential crosstalk between the actin and microtubule cytoskeletal structures during cell division.

**Figure 6.**
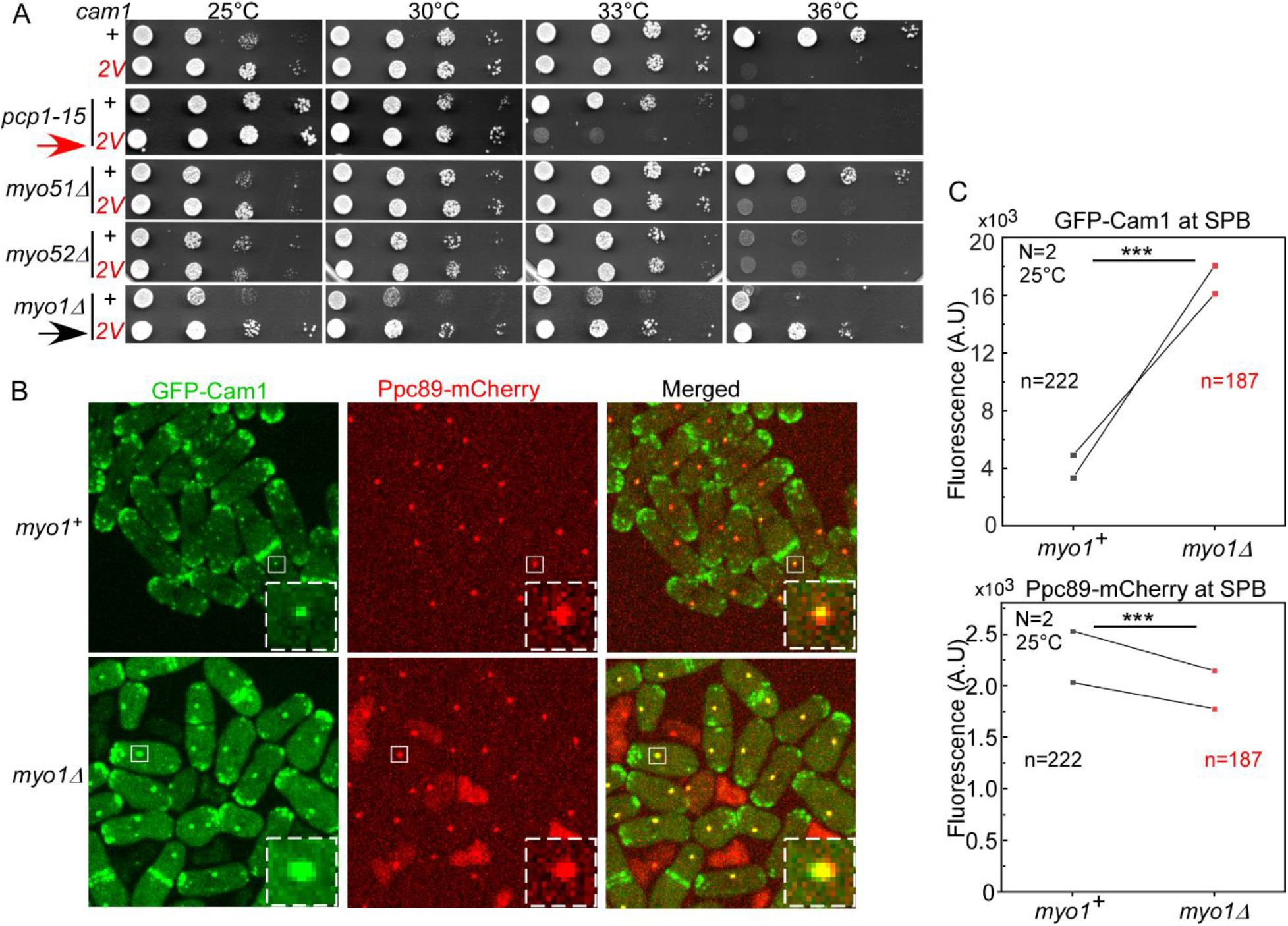
Deletion of myo1 rescues the temperature-sensitive growth of cam1-2V mutant. (A) Ten-fold dilution series of yeasts. *cam1-2V* showed a negative genetic interaction (red arrow) with *pcp1-15*. In comparison, *cam1-2V* showed no genetic interaction with either *myo51Δ* and *myo52Δ*, but it interacted positively (black arrow) with *myo1Δ*. (B) Micrographs of either *wild-type* (*myo1^+^*) or *myo1Δ* cells expressing both GFP-Cam1 (green) and Ppc89-mCherry (red). Dashed Box: magnified view of a SPB (box). (<u>C</u>) Paired dot plots of the fluorescence intensity of GFP-Cam1 at the SPBs. The number of GFP-Cam1 molecules at the SPBs increased by ∼3 folds in the *myo1Δ* cells, compared to the *wild-type* cells. Data is pooled from two independent biological repeats. ***: P<0.001, based on two-tailed student’s t tests.

## DISCUSSION

In this study, we discovered that the interaction between the cytoskeletal structures and calmodulin requires Ca^2+^ in the fission yeast *S. pombe*. Our finding provides a likely answer to the long-standing question of why fission yeast Cam1, unlike its budding yeast orthologue, requires Ca^2+^ for its essential functions (Moser et al., 1995). Ca^2+^-Cam1, together with the pericentrin-like Pcp1, may promote the duplication and assembly of the SPB. In contrast to Pcp1, the unconventional myosins interact with Cam1 with or without Ca^2+^ in the actin cytoskeletal structures. However, the functions of these myosins may be modulated by Ca^2+^-Cam1 during cytokinesis. The cytokinesis-related calcium influx could promote Cam1’s binding to Ca^2+^. Overall, Cam1 may act as a toggle between cytokinesis mediated by the unconventional myosins and mitosis promoted by Pcp1, dependent on the availability of Ca^2+^.

Ca^2+^-Cam1 shall be an integrative part of SPB throughout the cell cycle. This partially explains why the fission yeast calmodulin requires Ca^2+^ for its essential function. In contrast, the budding yeast cells dependent on a calmodulin mutant, that has completely lost its capacity to bind Ca^2+^, remain viable with no defects in either cell growth or division (Geiser et al., 1991). However, the similar fission yeast *cam1* mutation is lethal (Moser et al., 1995). Our inability to isolate the *cam1::cam1-3V* mutant so far concurs with this earlier finding. These Cam1 mutants that are unable to bind to Ca^2+^ are likely absent from the SPB completely, resulting in a failure in the SPB duplication and assembly. Therefore, one critical Ca^2+^-dependent role of Cam1 is to serve as a structural component of the microtubule-organizing center. Since the vertebrate calmodulin can replace Cam1 in fission yeast cells (Moser et al., 1995), this function of Cam1 could be conserved in vertebrate centrosomes.

Ca^2+^ shall promote both the direct interaction between Cam1 and Pcp1 and the activities of this pericentrin-like protein. This is supported by both microscopy and genetic data. When replacing the endogenous Cam1 with Cam1-2V, the majority of Pcp1 molecules are lost from SPBs even at the permissive temperature. The remaining fraction (30%) of Pcp1 could be retained due to its interaction with the other SPB protein, Ppc89 (Chen et al., 2024). Genetically, over-expression of Pcp1 alone is sufficient to rescue the temperature-sensitivity of *cam1-2V*. In contrary, the hypomorphic mutant *pcp1-15* exhibits strong negative genetic interaction with *cam1-2V*. Such a synergetic relationship between Ca^2+^-Cam1 and pericentrin may well be conserved evolutionarily.

The essential function of Pcp1-Cam1 complex is likely to promote the duplication and early assembly of the daughter SPB. During G1/S phase of the fission yeast cell cycle, the mother SPB assembles a half-bridge through the scaffold protein Sfi1 before the self-duplication (Bestul et al., 2017; Rüthnick et al., 2021). Both Cam1 and Pcp1 are some of the earliest components of the daughter SPB (Bestul et al., 2017). Therefore, the frequent loss of SPB, as well as the presence of a significant portion of either anucleate cells, in the *cam1-2V* mutant, is likely due to either failed duplication or early assembly of the daughter SPB. Except for Pcp1, the loss of Cam1 from the SPB has only a minor effect on the assembly of other SPB proteins that we examined, including Ppc89, Cdc11, the gamma-tubulin complex, and Plo1 kinase. Consistent with this result, most mitotic spindles, nucleated by the gamma-tubulin complex, extend normally in the *cam1-2V* mutant cells. Therefore, the critical function of Pcp1-Cam1 at the SPB is to promote the daughter SPB duplication and assembly.

Unlike Pcp1, the unconventional myosins bind to Cam1, regardless of its Ca^2+^ binding status, at the actin cytoskeletal structures. This is supported by the normal localization of Cam1-2V and -3V in the cell tips, where there are both actin patches and cables. This implies that these Cam1 mutants continue to interact with the type I and V myosins. This is also consistent with our observation that both Cam1 mutants remain at the division plane, which needs their interaction with type V myosins Myo51 and Myo52. Our data suggests that Cam1 continues to bind to these unconventional myosins even without requiring Ca^2+^, like the budding yeast calmodulin (S. E. Brockerhoff & Davis, 1992). Our in vivo result is in line with the in vitro observation that calmodulin binds the IQ motif of these unconventional myosins independent of Ca^2+^ (Perreault-Micale et al., 2000).

Ca^2+^-Cam1 likely modulates the motor activities of the two unconventional type V myosins. Both type V myosins promote the assembly of the actomyosin contractile ring and the ring constriction (Laplante et al., 2015). The accelerated contractile ring assembly and constriction in the *cam1* mutant cells suggest an elevated activity of Myo51 and Myo52. Combined, our results suggest that Ca^2+^-Cam1 plays a critical role in cytokinesis through modulating the activities of type V myosins.

The two novel suppressors of *cam1-2V* mutant suggest that Cam1 has a dual role in both mitosis and cytokinesis. One suppressor is the gain of function mutant of *pcp1*. The over-expression of Pcp1 likely promotes SPB duplication in the *cam1* mutant cells at the restrictive temperature. Our finding echoes the earlier finding in the budding yeast in which the *pcp1* orthologue *spc110* is a high-copy suppressor of the calmodulin mutant *cmd1-1* (Sundberg et al., 1996). The other suppressor, a surprising one, is the loss of function mutant of *myo1*. The deletion of *myo1* increases the localization of Cam1 at the SPB. This may indirectly enhance the activity of Cam1 in the SPB. Therefore, the essential functions of Ca^2+^-Cam1 are likely to be two-fold, in both the cell division dependent on the SPB and modulating cytokinesis requiring unconventional myosin.

Cam1-Ca^2+^ may be promoted by the calcium transients associated with fission yeast cell division. Dividing fission yeast cells trigger two separate cytokinetic calcium spikes (Poddar et al., 2021). One of them, the constriction spike, occurs during M phase when the ring starts to constrict. The other, the separation spike, appears at the end of daughter cell separation. These calcium transients may come from the influx through mechanosensitive calcium channels such as Pkd2 (Poddar et al., 2022). Thus, the intracellular calcium transients may promote the formation of the Pcp1-Cam1 complex at the SPB. Alternatively, the other potential source of Ca^2+^ is the ER, which stores high concentration of Ca^2+^ and is in close proximity to the SPB.

Overall, our results demonstrate that Ca^2+^-Cam1 serves as both an essential constitutive component of the SPBs and the actin cytoskeletal structures during cell division. It plays paradoxical roles in mitosis and cytokinesis.

## MATERIAL AND METHODS

### Fission Yeast Genetics

Fission yeast was cultured by following the standard procedures (Moreno et al., 1991). We used a SporePlay+ tetrad microscope (Singer, England) to dissect the tetrads. To prepare a ten-fold dilution series, the yeasts were first inoculated overnight in liquid YE5s medium in a 50 ml flask in a water bath shaker at 25°C. The next morning, the confluent cell cultures were diluted ∼20-folds and allowed to grow for another 5–6 hours before preparing a series of ten-fold dilutions of the yeast cultures, which were then deposited onto the agar plates. The yeast cultures were incubated at designated temperatures for 2-3 days. The plates were scanned with a photo scanner (Epson, USA). All the yeast strains used in this study have been listed in Supplemental Table 1.

### Construction of *cam1* mutants

To express sfGFP-Cam1 and -3V driven by the actin promoter exogenously, we amplified the ORF of both the *wild-type cam1* and *cam1-3V* from plasmids V232 and V240 (gifts from Trisha Davis), respectively, through PCR using primers P1082 and P1084. Both were subcloned into the vector pAV-0714 (a gift from Sophie Martin) using HiFi assembly (NEB, USA). The resulting plasmids V298 and V299 were confirmed through Sanger Sequencing. The coding sequence of Cam1-3V carried three missense mutations, E33V (G<u>A</u>A to G<u>T</u>A), E69V (G<u>A</u>A to G<u>T</u>A), and E106V (GAG to G<u>TC</u>). The sfGFP-Cam1-2V construct was generated through PCR-based site-directed mutagenesis using Q5 DNA polymerase (NEB). We reverted the E33V mutation in the Cam1-3V coding sequence using primers P1673 and P1674. Positive clones were selected through PCR and confirmed through Sanger sequencing. 1 µg of the plasmids was linearized by digestion with the restriction enzyme Afe1 (NEB). The linearized plasmids were purified and transformed into yeast via the Li-Acetate method. Positive transformants were selected on an EMM-ura plate and confirmed through PCR using primers P1092 and P829.

To construct the *cam1::cam1-2V and 3V* mutants, we used the two-step marker reconstitution method (Tang et al., 2011). First, the DNA fragment encoding the C-terminally truncated His5 (*his5CterΔ*) and *ura4* from the pH5C vector (Tang et al., 2011) was amplified by PCR using primers P830 and P831. The primers included the sequences homologous to the 3′ untranslated region (UTR) of *cam1*. The amplified DNA fragment was transformed into wild-type yeast (Y986). The resulting yeast strain Y1403 was confirmed through PCR and Sanger sequencing. Next, the coding sequence for *cam1-3V* including 3′ UTR was amplified from the plasmid V240 (a gift from the Trisha Davis lab) (Moser et al., 1995). It was then cloned into the vector pH5D (containing His5Cter) (Tang et al., 2011). The recombinant vector QC-V245 was used as a template to amplify the sequence of *cam1-3V-*His5Cter. The PCR product was transformed into Y1403 to replace the endogenous *cam1* gene with the *cam1-3V* mutant through homologous recombination. The transformants were selected on EMM-his plates and were further screened for temperature-sensitive mutants on plates supplemented with Phloxine B at 36°C for 2 days. The *cam1* locus of the temperature-sensitive clones was amplified using primers P1140 and P600 and sequenced through Sanger sequencing. Out of 194 transformants that we screened, we found only 5 temperature-sensitive mutants. Among them, only two carried the mutant allele *cam1::cam1-2V,* and none carried the allele *cam1::cam1-3V*.

### Immunoblots

To determine the expression level of sfGFP-Cam1, 2V, and 3V, respectively, 30 O.D. of yeast cells were harvested by centrifugation at 1,500 g for 5 minutes. The pellet was washed once with 1 mL of sterile water. The pelleted cells were re-suspended with 300μL of lysis buffer (50 mM Tris-HCl pH 7.5, 100 mM KCl, 3 mM MgCl2, 1 mM EDTA, 1 mM DTT, 0.1% Triton X-100, and protease inhibitors (Halt protease inhibitor cocktail; Thermo Fisher #1862209)). The cells were mechanically homogenized with 300mg of glass beads using a bead beater (BeadBug, Benchmark Scientific, USA). They were lysed for five cycles of 1 min disruption plus 1 min incubation on ice. The cell lysis was immediately mixed with preheated 5x Laemelli sample buffer before being heated for 10min at 100°C. After being centrifuged at 15,000 RPM for 1min, 200µl of supernatant was collected and 20µl was used for SDS-PAGE gel electrophoresis (Mini-PROTEIN TGX 10% precast, BioRad #4561033, USA). The gels were either stained with Brilliant blue R-250 (FisherBiotech cat# 6104-59-2) or transferred to PVDF membrane (Amersham, #10600023) for immunoblots. The blots were first incubated with the anti-GFP antibodies (1:1000; #11814460001; Sigma-Roche) overnight at 4°C, followed by horseradish peroxidase–conjugated secondary antibodies (1:10,000; #sc-516102; Santa Cruz) for 1h at the room temperature. The blots were developed with chemiluminescent reagents (Pierce ECL Western Blotting Substrate, ThermoFisher Scientific, #32209, USA). We quantified the immunoblots using NIH ImageJ. To quantify the density of a band, we drew a rectangle around it to measure the average intensity. This was then subtracted with the background intensity, measured in an area where there were no visible bands.

### Purification of recombinant Cam1 and Cam1-3V

The cDNA sequences of *S. pombe* Cam1 and the 3V mutant were both cloned into the modified pQE vector V290 (a gift from Steven Chou) to the C-terminus of GST and a 17aa linker (SDLVPRGSLEVLFQGPGG) that includes a thrombin protease digestion site through Hi-Fi assembly (NEB, USA). The resulting plasmids, V399 and V409, were confirmed through Sanger sequencing.

Both GST-tagged Cam1 and Cam1-3V proteins were expressed in *E. coli* BL21 (DE3) cells (NEB, cat#C2527) and purified by affinity chromatography using glutathione–Sepharose resin (ThermoFisher, cat#16100). Briefly, the bacteria were inoculated in LB broth supplemented with 100 μg/ml ampicillin (RPI, cat# A40040) overnight in an orbital shaker at 37°C. The overnight culture was diluted 1:1000 into 4L of LB medium and grown to mid-log phase. The culture was then induced with 0.5 mM isopropyl β-d-thiogalactopyranoside (IPTG) (RPI, cat#367-93-1) at 37 °C for 4 hrs. The bacterial culture was harvested by centrifugation at 3,500g for 10 mins at 4°C (Beckman, Avanti centrifuge J-25I). The pellet was resuspended in GST column buffer (50mM Tris HCl pH 8.0, 150mM NaCl, 1mM DTT), supplemented with 1mM PMSF and 1 tablet of Protease inhibitor cocktail (Roche; Cat#11836170001), and were sonicated (U.S. Solid Ultrasonic Homogenizer) for 10 cycles of short pulses of 45s on and 45s off. The cell lysate was centrifuged at 20,000g for 30 mins at 4°C (Beckman, Avanti centrifuge J-25I). The supernatant was mixed with 1ml glutathione beads for 2h at 4°C on a rocking platform. The lysate and resin were poured into a disposable column (Bio-Rad, cat#731-1550) and allowed to pack slowly through gravity. The flowthrough was collected and the resin was washed with 10 column volumes of GST column buffer. The protein was eluted with 5ml elution buffer (column buffer plus 10mM reduced glutathione (Thermo Fisher, Cat#78259)). The recombinant proteins were detected through electrophoresis in a SDS-PAGE gel. The fractions containing the recombinant proteins were pooled and dialyzed overnight against 2L buffer (50mM Tris HCl pH 8.0, 150mM NaCl, 1mM DTT). Aliquots of concentrated protein were flash-frozen in liquid nitrogen for long-term storage at −80 °C. The concentrations of recombinant protein were measured using the Pierce BCA Protein Assay Kit (Thermo Fisher Scientific, cat#23227) on a plate reader (SpectraMax M5).

The purified recombinant GST-Cam1 and GST-Cam1-3V were mixed with either CaCl_2_ (4mM) or EDTA (2mM) in 25µl binding buffer (50mM Tris HCl pH 8.0, 150mM NaCl, and 1mM DTT) and incubated for 20 mins on a rocking platform at the room temperature. 5X Lamelli sample buffer was added to each reaction and heated at 90°C for 5 mins. The samples were then analyzed using 15% SDS–PAGE gel. The gels were stained with Brilliant blue R-250 (Fisher Biotech, cat#BP101-25) and destained before being scanned with a scanner (Epson perfection V550 photo).

### Microscopy

For microscopy, the yeast cells were inoculated in liquid YE5s media at 25°C for two days before being harvested during the exponential growth phase at a density between 0.5 and 1x10^6^ cells/ml. For temperature-sensitive mutants, exponentially growing cultures were inoculated at the restrictive temperature of 36°C for 3-4 hrs before experiments. The cells were resuspended in 50 μl of YE5S and 6 μl of the resuspended cells were spotted onto a gelatin pad (25% gelatin in YE5S) on a glass slide. The cells were sealed under a coverslip (#1.5) with VALAP (1:1:1 mixture of Vaseline, lanolin and paraffin) before imaging.

Live microscopy was carried out on a spinning disk confocal microscope equipped with an EM-CCD camera. The Olympus IX71 microscope was equipped with objective lenses of 60× (NA = 1.40, oil) and 100× (NA = 1.40, oil), a motorized Piezo Z Top plane (Applied Scientific Instrumentation, Eugene, OR, USA) and a confocal spinning disk unit (Confocal Scanner Unit -X1, Yokogawa, Tokyo, Japan). The images were captured on an Ixon-897 EMCCD camera (Andor, USA). Solid-state lasers of 488 and 561 nm were used. Unless specified, the cells were imaged with a Z-series of 15 slices at a step size of 0.5 μm. Time-lapse microscopy was carried out in a dedicated dark room, maintained at ∼23°C. To minimize temperature variations among the experiments, we always imaged the wild-type and mutant cells on either the same or consecutive days

For time-lapse microscopy in a liquid media, 20–40 μl of exponentially growing cell culture was spotted onto a glass coverslip (#1.5) at the bottom of a 10 cm Petri dish (Cellvis, USA). The dish had been coated with 50μl of 50μg/ml lectin (Sigma, L2380) and allowed to dry overnight at 4°C. The cells were allowed to attach to the glass slip for 10 minutes before 2ml of YE5s media was added. For experiments at the restrictive temperature, we used a temperature-controlled chamber with an inserted temperature sensor for a 10cm petri dish (872-1928, OkaLab) set at sample-controlled mode. The temperature was measured using a thermometer submerged in the media. The temperature usually varied by 0.15°C throughout the time-lapse movies.

For visualizing the fission yeast nuclei, we fixed the cells with 70% ethanol and stained them with 1ug/ml DAPI (Roche #1023627001). To visualize the cell wall, we fixed the yeast cells with with 4% paraformaldehyde, prepared from 16% fresh stock solution (Electron Microscopy Science, USA). The fixed cells were washed with TEMK buffer (50 mM Tris-HCL pH 7.4, 1 mM MgCl2, 50 mM KCl, 1 mm EGTA, pH 8.0). They were then stained with 1ug/ml calcofluor (1mg/mL Sigma #18909) on a rocking platform for 10 min at room temperature in the dark. The stained cells were pelleted and resuspended in 50 μL TEMK, 6μL of which was spotted on a glass slide directly and sealed under a coverslip (#1.5) with VALAP. We used an Olympus IX81 microscope equipped with a CCD camera and a mercury lamp to image these fixed cells.

### Microscopy image Processing and Analysis

We used NIH ImageJ, installed with either freely available or customized macros and plug-ins, to process all the microscopy data. For quantitative analysis, the fluorescence micrographs were first corrected both for X–Y drift by using StackReg (Thevenaz et al., 1998) and for photo-bleaching by using EMBL Tools (Rietdorf, EMBL Heidelberg). For most quantitative fluorescence measurements, the average fluorescence intensities of all the Z-slices were used.

Localization to SPBs of a protein was measured by counting the average fluorescence intensity within a 0.8μm (8 pixels)-diameter circle for 100X and (5 pixels) 60X lenses, respectively. It was subtracted from the fluorescence intensity in a ring of 1.25 μm (12 pixels) in diameter, concentric with the former. For the latter, a ring of 1.5 µm (9 pixels) in diameter was used instead. The division plane localization of a protein was quantified by measuring the fluorescence intensities within a 4.35 by 1 μm (25 × 6 pixels) rectangle centered on the equator. It was subtracted by the average fluorescence intensity in a 4.35 by 0.7 μm (25 X 4 pixels) rectangle adjoined to the equatorial plane.

The ImageJ Plug-in Cell Counter was used to count cells with various phenotypes. We counted the lysed daughter cells throughout the time-lapse movies using NIH ImageJ. We counted all cell separation events over a 2-hr period. Similarly, we counted the lysed cells during cytokinesis throughout 2-hour window, before calculating the percentage of lysed cells.

## Acknowledgment

The authors would like to thank the members of the Chen lab at the University of Toledo for providing technical assistances. The authors would like to thank Trisha Davis (University of Washington), Kathy Gould (Vanderbilt University), Steven Chou (University of Connecticut Health Science Center), Vladimir Sirotkin (SUNY Upstate Medical Center), and National Bio-Resource Project (NBRP, Japan) for sharing yeast strains and plasmids. This work has been supported by the grants NSF 2144701 and NIH R01GM144652 to QC.

## Supplemental Materials

**Table 1.**
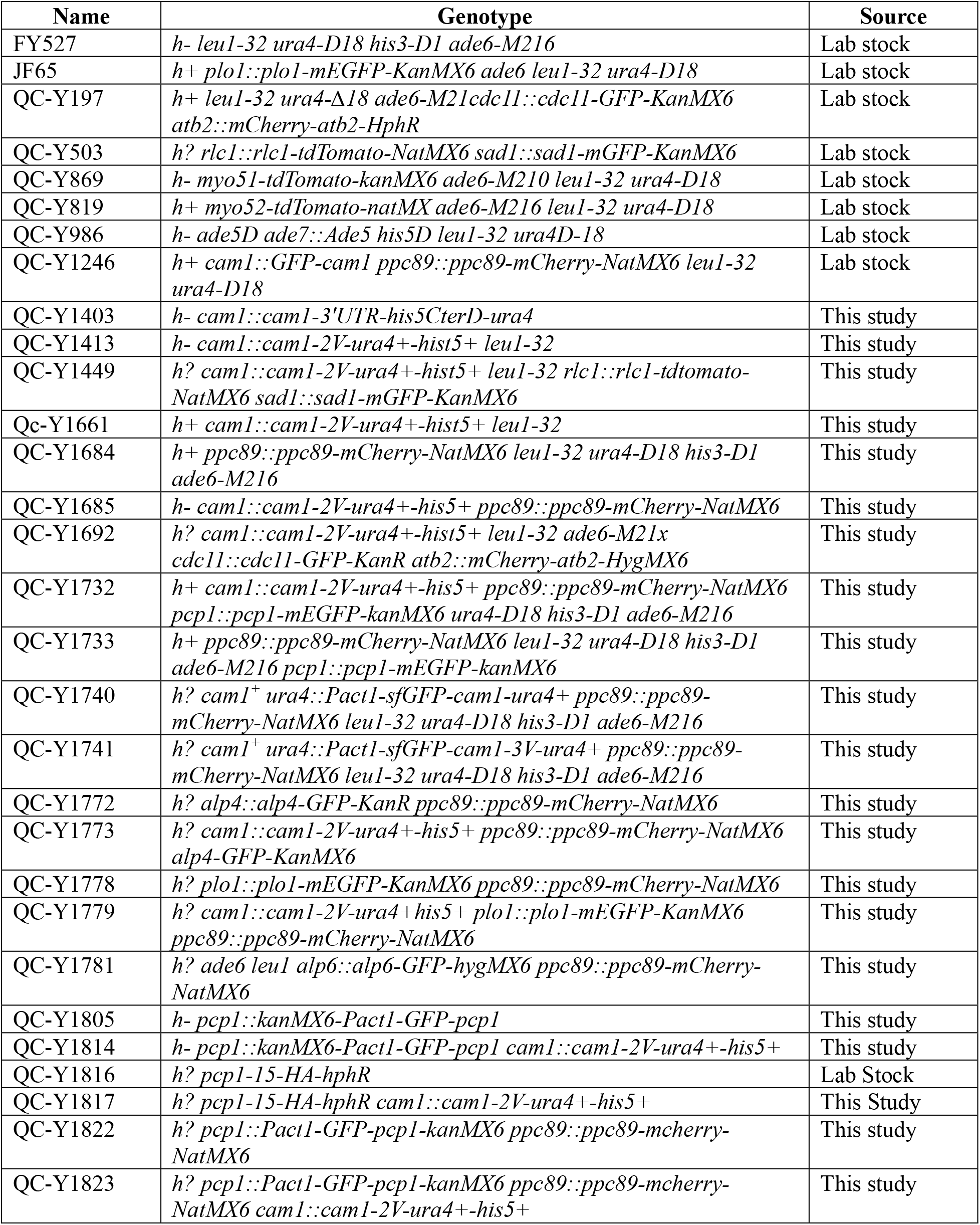
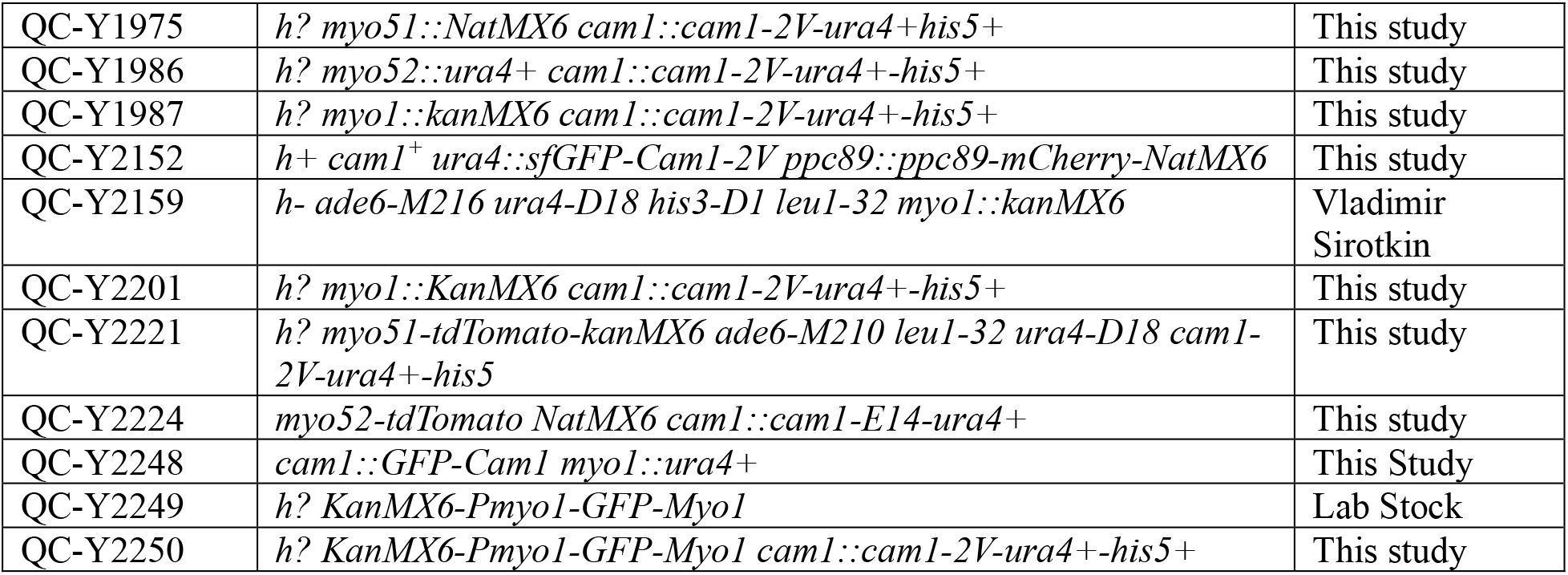
Yeast strains used in this study.

**Supplemental Figure S1.**
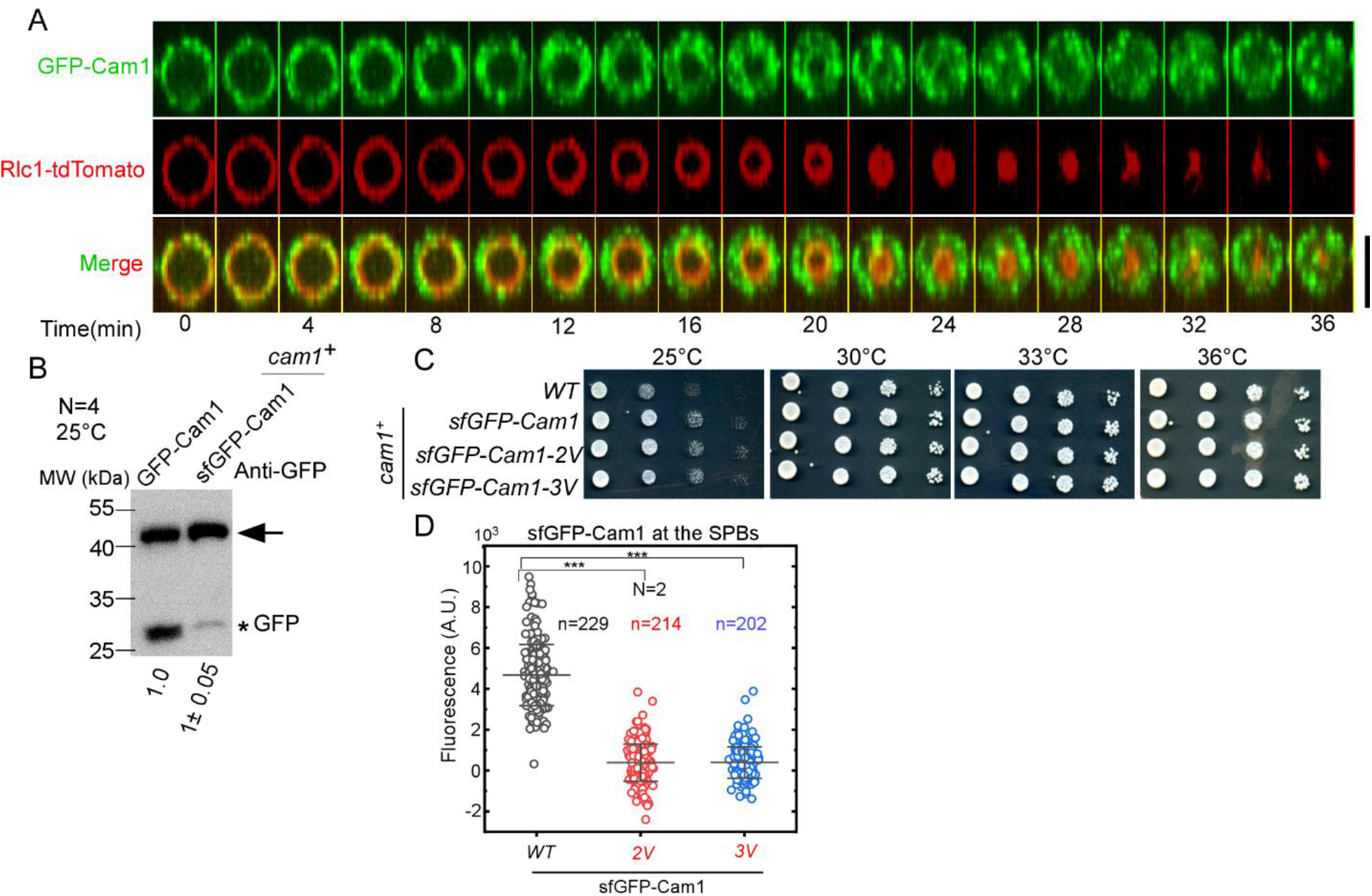
*2V* or *3V* mutations of Cam1 reduces its localization at the SPB. (A) Time-series of the division plane (cross-section view shown) of a cell co-expressing GFP-Cam1 (green) and Rlc1-tdTomato (red). GFP-Cam1 was localized to numerous puncta in a donut-shaped disk at the equatorial division plane. Number: time in minutes after the start of the contractile ring constriction. (B) Scanned anti-GFP blots of the whole cell lysates of the yeasts expressing either endogenous GFP-Cam1 or exogenous sfGFP-Cam1 driven by the actin promoter. (C) Ten-fold dilution series of yeasts. (D) Dot plot of the fluorescence intensities of sfGFP-Cam1 (WT), -2V and 3V at the SPBs of asynchronized cells. Cam1-2V and -3V were largely absent from the SPBs even at the permissive temperature of 25°C. Data is pooled from two independent biological repeats. ***: P<0.001, based on two-tailed student’s t tests.

**Supplemental Figure S2.**
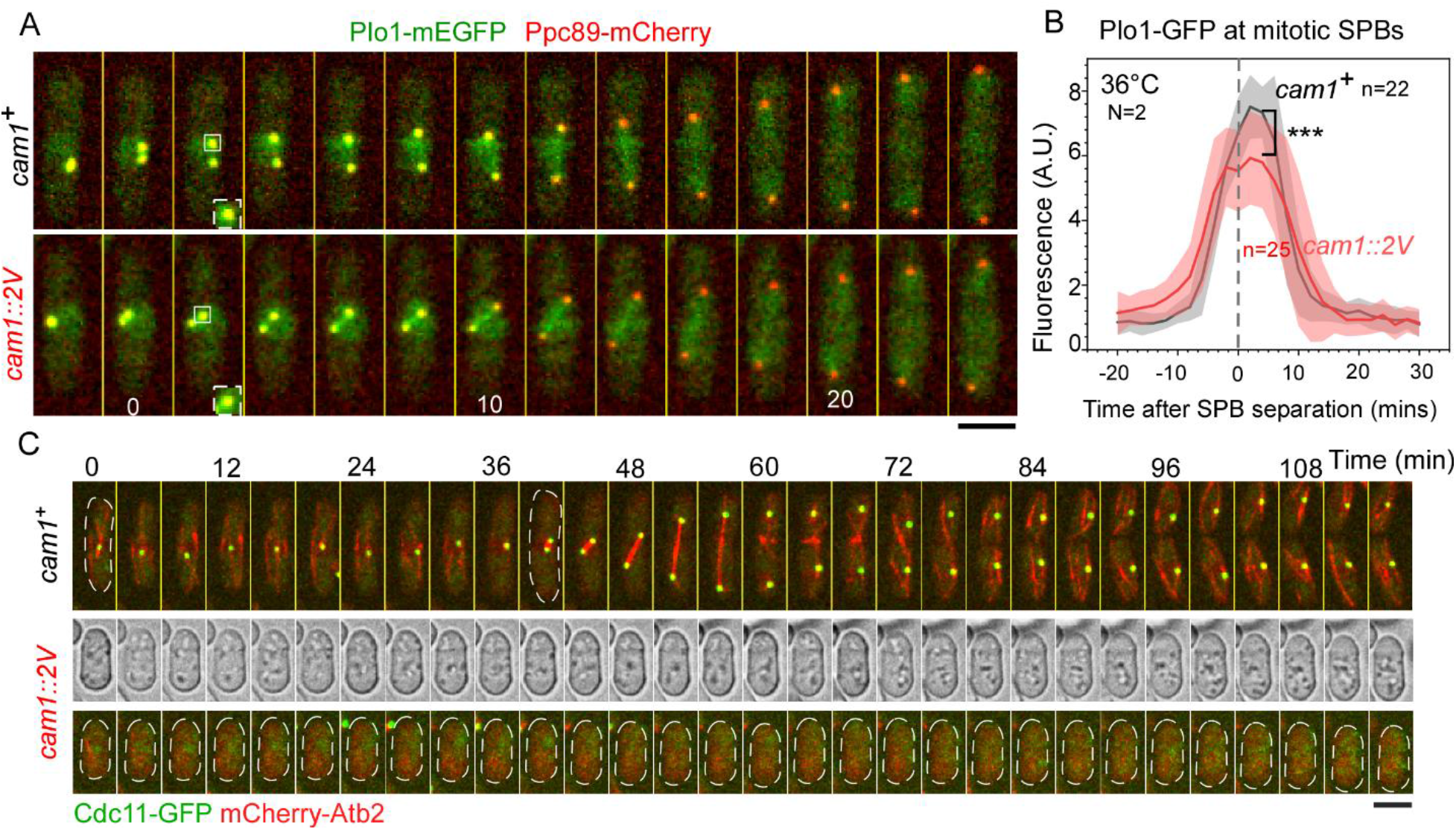
The localization of Plo1-GFP at the mitotic SPB of *cam1-2V* cells decreases by *cam1-2V* at the restrictive temperature. (A) Time series of a *wild-type (cam1^+^)* and *cam1-2V* cells co-expressing both Plo1-GFP (green) and Ppc89-mCherry (red) at 36°C. (B) Average time courses of the fluorescence intensities of Plo1-GFP at the SPBs of either the *wild-type* (*cam1^+^*) or *cam1-2V* cells. Cloud: standard deviation. (C) Time series of a *wild-type* (top) and SPB-less *cam1-2V* (bottom) cell co-expressing Cdc11-GFP (SPB marker, green) and mCherry-Atb2 (microtubule marker, red) at 36°C. No mitotic spindles appeared in the mutant cell in the two-hour window. In contrast, the *wild-type* cell assembled and extended mitotic spindles within this timeframe. Number: time from the beginning of this time-series in minute. ***: P<0.001, based on two-tailed student’s tests. Bar: 5µm.

**Supplemental Figure S3.**
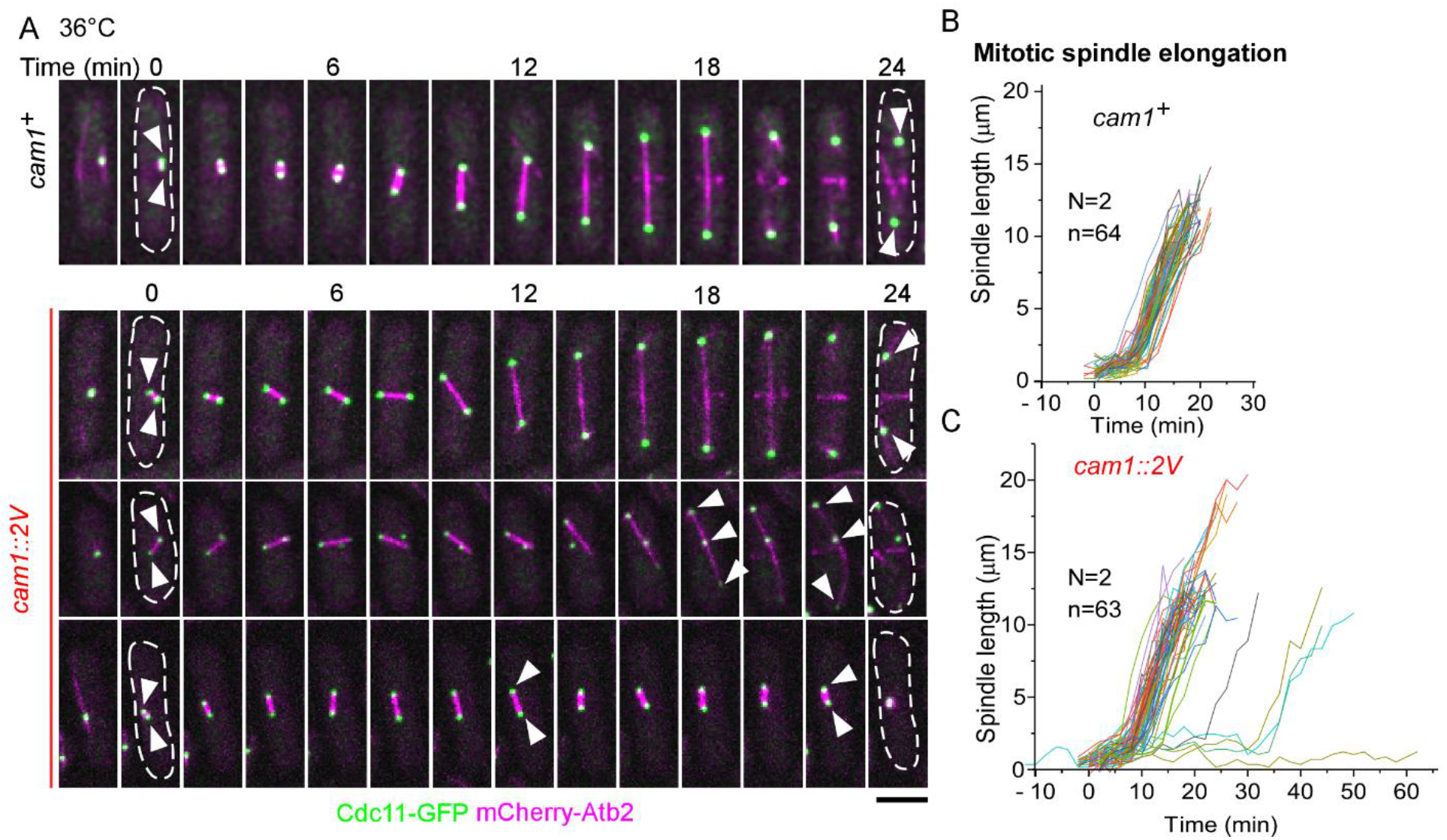
The mitotic spindle of most *cam1-2V* mutant cells elongates normally. (A) Time series of a *wild-type* (top) and three representative *cam1-2V* dividing cells (bottom) expressing both Cdc11-GFP (green) and mCherry-Atb2 (magenta), respectively. Number: Time in minutes after the SPB separation. Arrowhead: SPBs. (B-C) Line plots of the time courses of the length of the mitotic spindles in either *wild-type* (*cam1^+^*, B) or mutant (*cam1::2V,* C) cells. Data is pooled from two independent biological repeats. Bar: 5µm.

**Supplemental Figure S4.**
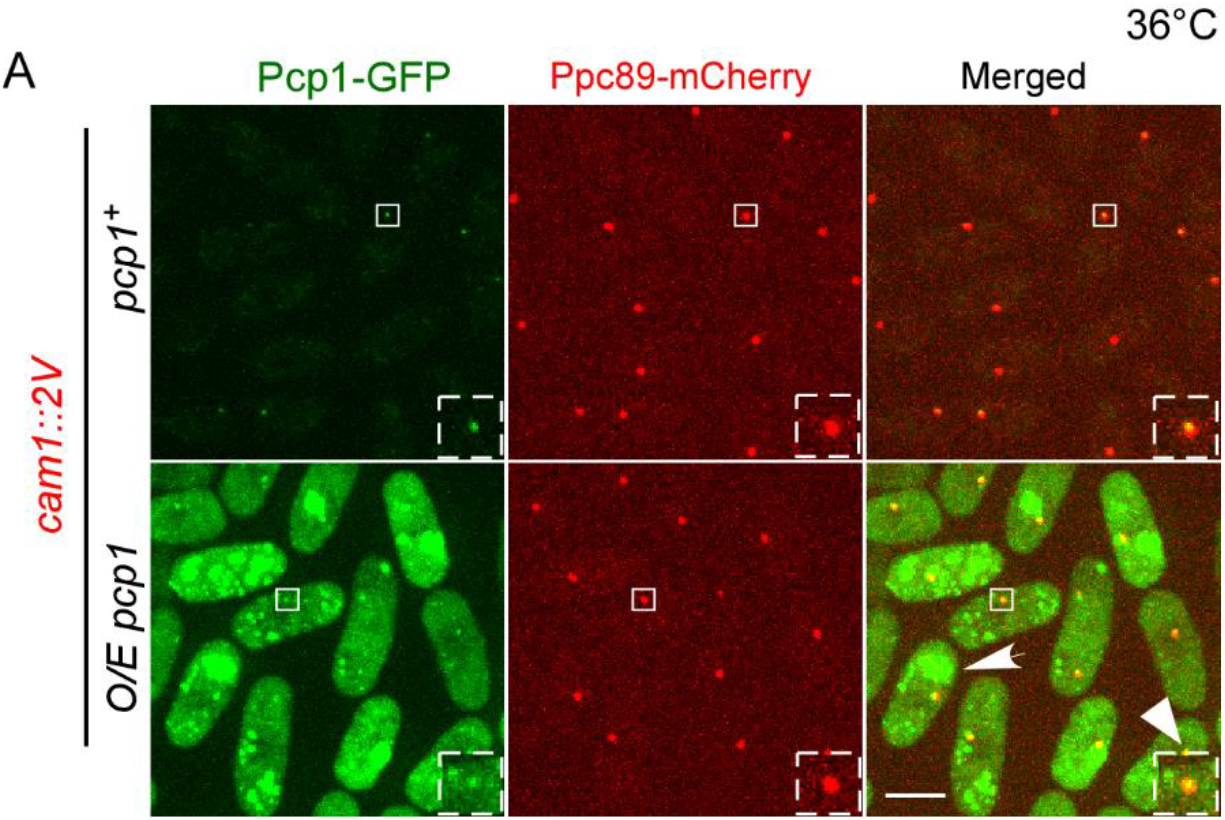
Over-expressed GFP-Pcp1 diffuses throughout the cytoplasm of *cam1-2V* cells. Representative micrographs of *cam1-2V* mutant cells expressing both Ppc89-mCherry (SPB marker, red) and either endogenous Pcp1-GFP (green, *pcp1^+^*) or over-expressed GFP-Pcp1 (*O/E pcp1*) at 36°C. Dashed Box: magnified view of representative SPBs (box). Unlike the endogenously expressed Pcp1, the over-expressed Pcp1 diffused throughout the cytoplasm except for a few puncta. Representative data from three independent biological repeats is shown. Bar: 5µm.

**Supplemental Figure S5.**
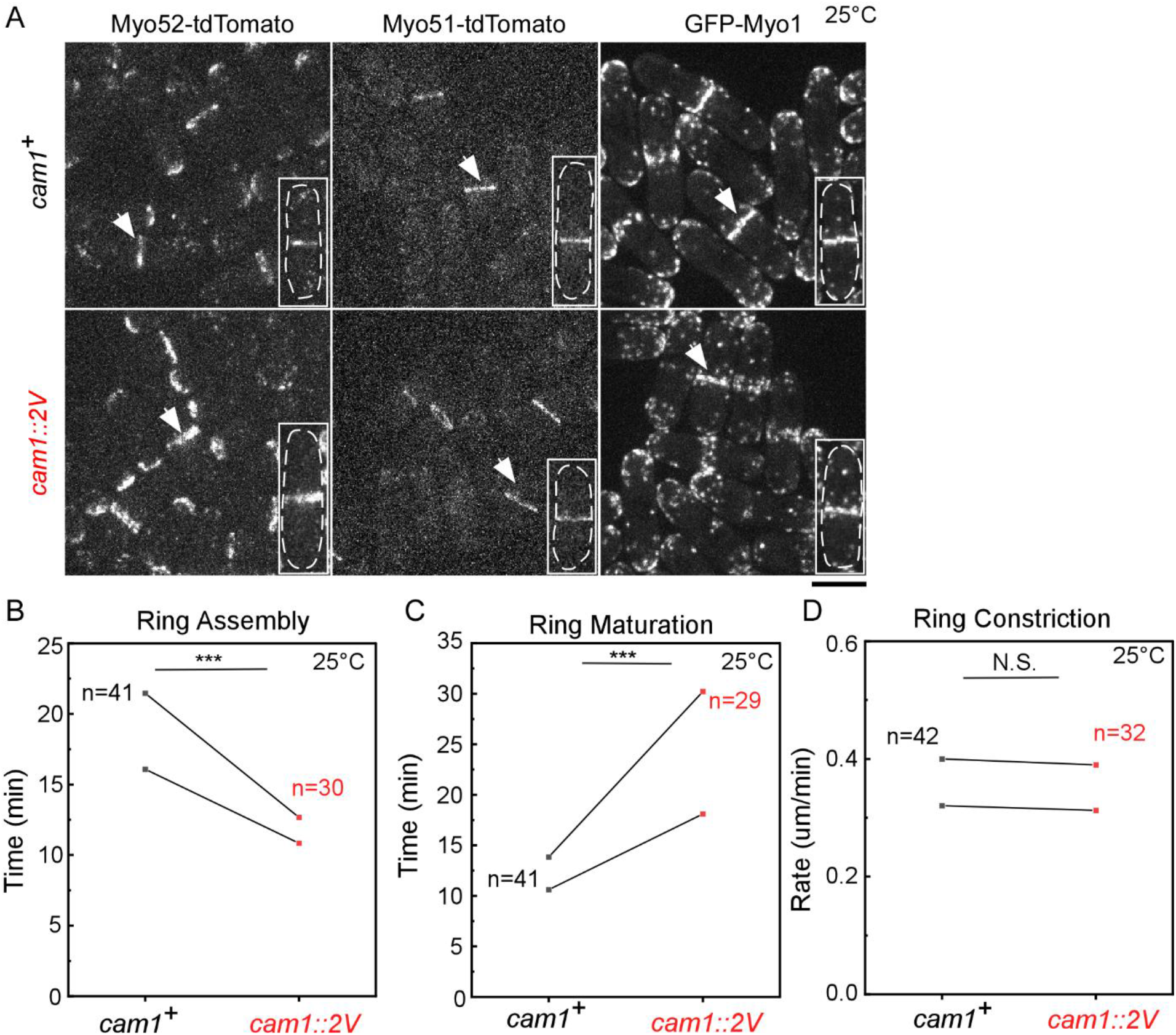
Cytokinesis in the *cam1-2V* cells is unchanged from the *wild-type* cells at the permissive temperature. (A) Micrographs of the *wild-type* (top) and *cam1-2V* (bottom) cells expressing either Myo52-tdTomato, Myo51-tdTomato, or GFP-Myo1 at 25°C. Box: magnified view of a representative dividing cell. Arrow: the equatorial division plane. (B-D) Paired dot plots of the durations of the contractile ring assembly (B) and maturation (C), in addition to the ring constriction rate (D) of the *wild-type* (*cam1^+^*) and *cam1-2V* cells at 25°C.

**Supplemental Figure S6.**
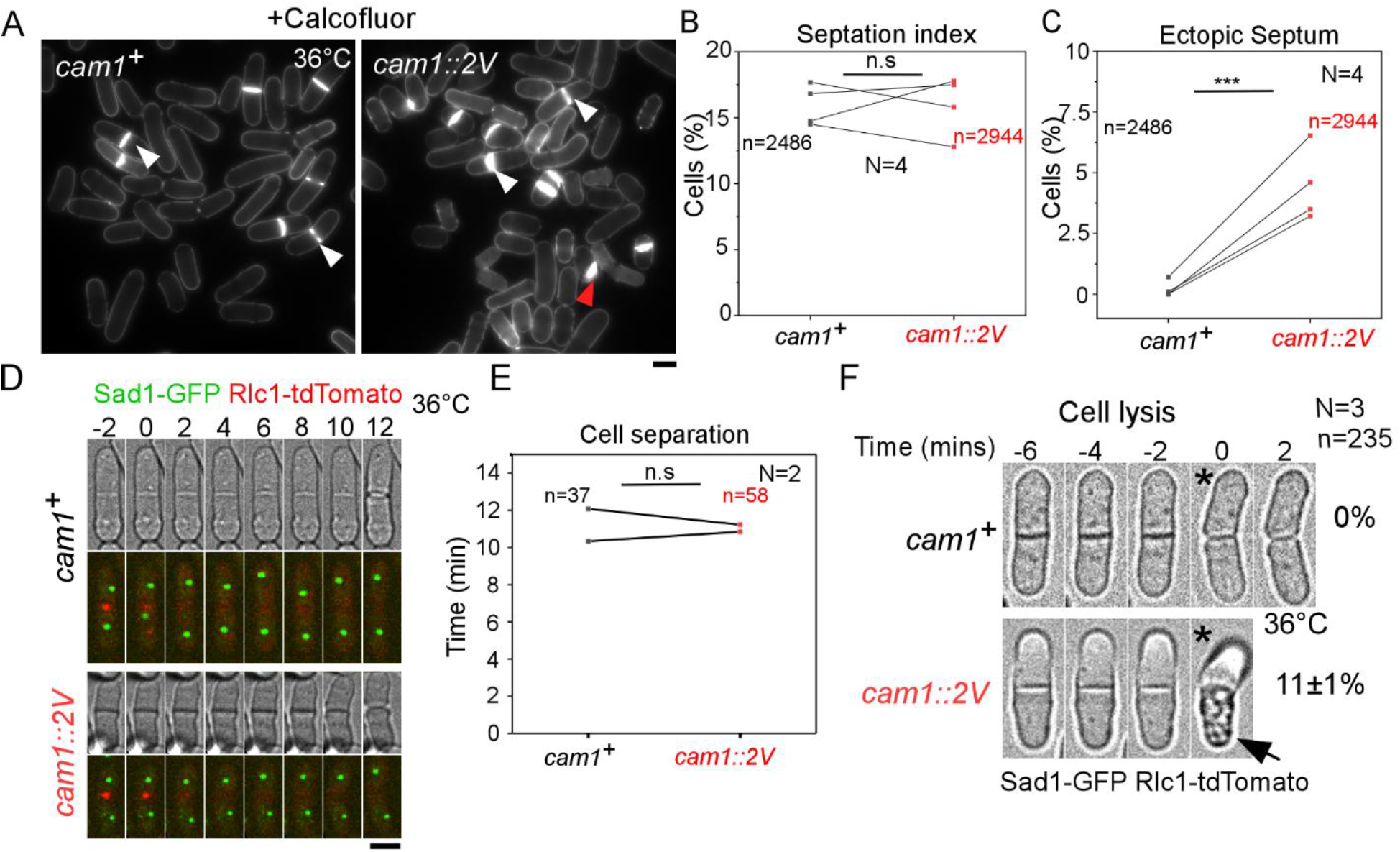
*cam1-2V* cells lyse frequently after the daughter cell separation at the restrictive temperature. (A) Micrographs of calcofluor-stained *wild-type* (left) and *cam1-2V* (right) cells at 36°C. White arrowhead: normal septum. Red arrowhead: ectopically deposited septum. (B-C) Paired dot plots of the fraction of the cells with either septum (B) or ectopic septum (C) at 36°C. About 5% of the mutant cells erroneously deposited septum at either the cell tip or side. (D) Time series of a representative *wild-type* (top) and *cam1-2V* (bottom) dividing cell, both expressing Sad1-GFP (green) and Rlc1-tdTomato (red). Number: Time in minute after the ring closure. (E) Paired dot plot of the duration of the cell separation. (F) Time series of a representative *wild-type* (top) and *cam1-2V* (bottom) separating cell. About 11±1% (average ± standard deviation) of the mutant cells lysed (arrow) immediately after the separation. Number: Time in minutes after daughter cell separation (asterisk). N.S.: P>0.05. ***: P<0.001, based on two-tailed student’s t-tests.

## References

Babu, Y. S., Sack, J. S., Greenhough, T. J., Bugg, C. E., Means, A. R., & Cook, W. J. (1985). Three-dimensional structure of calmodulin. Nature, 315(6014), 37–40. 10.1038/315037a0

Bestul, A. J., Yu, Z., Unruh, J. R., & Jaspersen, S. L. (2017a). Molecular model of fission yeast centrosome assembly determined by superresolution imaging. Journal of Cell Biology, 216(8), 2409–2424. 10.1083/jcb.201701041

Bestul, A. J., Yu, Z., Unruh, J. R., & Jaspersen, S. L. (2017b). Molecular model of fission yeast centrosome assembly determined by superresolution imaging. Journal of Cell Biology, 216(8), 2409–2424. 10.1083/jcb.201701041

Brockerhoff, S. E., & Davis, T. N. (1992). Calmodulin concentrates at regions of cell growth in Saccharomyces cerevisiae. Journal of Cell Biology, 118(3), 619–629. 10.1083/jcb.118.3.619

Brockerhoff, S., Stevens, R., & Davis, T. (1994). The unconventional myosin, Myo2p, is a calmodulin target at sites of cell growth in Saccharomyces cerevisiae. The Journal of Cell Biology, 124(3), 315–323. 10.1083/jcb.124.3.315

Chen, J.-S., Igarashi, M. G., Ren, L., Hanna, S. M., Turner, L. A., McDonald, N. A., Beckley, J. R., Willet, A. H., & Gould, K. L. (2024). The core spindle pole body scaffold Ppc89 links the pericentrin orthologue Pcp1 to the fission yeast spindle pole body via an evolutionarily conserved interface. Molecular Biology of the Cell, 35(8), ar112. 10.1091/mbc.E24-05-0220

Cheung, W. Y. (1970). Cyclic 3′,5′-nucleotide phosphodiesterase. Biochemical and Biophysical Research Communications, 38(3), 533–538. 10.1016/0006-291X(70)90747-3

Crotti, L., Johnson, C. N., Graf, E., De Ferrari, G. M., Cuneo, B. F., Ovadia, M., Papagiannis, J., Feldkamp, M. D., Rathi, S. G., Kunic, J. D., Pedrazzini, M., Wieland, T., Lichtner, P., Beckmann, B.-M., Clark, T., Shaffer, C., Benson, D. W., Kääb, S., Meitinger, T., … George, A. L. (2013). Calmodulin Mutations Associated With Recurrent Cardiac Arrest in Infants. Circulation, 127(9), 1009–1017. 10.1161/CIRCULATIONAHA.112.001216

Crotti, L., Spazzolini, C., Tester, D. J., Ghidoni, A., Baruteau, A.-E., Beckmann, B.-M., Behr, E. R., Bennett, J. S., Bezzina, C. R., Bhuiyan, Z. A., Celiker, A., Cerrone, M., Dagradi, F., De Ferrari, G. M., Etheridge, S. P., Fatah, M., Garcia-Pavia, P., Al-Ghamdi, S., Hamilton, R. M., … Schwartz, P. J. (2019). Calmodulin mutations and life-threatening cardiac arrhythmias: Insights from the International Calmodulinopathy Registry. European Heart Journal, 40(35), 2964–2975. 10.1093/eurheartj/ehz311

Davis, T. N., Urdea, M. S., Masiarz, F. R., & Thorner, J. (1986). Isolation of the yeast calmodulin gene: Calmodulin is an essential protein. Cell, 47(3), 423–431. 10.1016/0092-8674(86)90599-4

Eng, K., Naqvi, N. I., Wong, K. C. Y., & Balasubramanian, M. K. (1998). Rng2p, a protein required for cytokinesis in fission yeast, is a component of the actomyosin ring and the spindle pole body. Current Biology, 8(11), 611–621. 10.1016/S0960-9822(98)70248-9

Flory, M. R., Morphew, M., Joseph, J. D., Means, A. R., & Davis, T. N. (2002). Pcp1p, an Spc110p-related calmodulin target at the centrosome of the fission yeast Schizosaccharomyces pombe. Cell Growth & Differentiation: The Molecular Biology Journal of the American Association for Cancer Research, 13(2), 47–58.

Fong, C. S., Sato, M., & Toda, T. (2010). Fission yeast Pcp1 links polo kinase-mediated mitotic entry to γ-tubulin-dependent spindle formation. The EMBO Journal, 29(1), 120–130. 10.1038/emboj.2009.331

Geiser, J. R., Sundberg, H. A., Chang, B. H., Muller, E. G., & Davis, T. N. (1993). The essential mitotic target of calmodulin is the 110-kilodalton component of the spindle pole body in Saccharomyces cerevisiae. Molecular and Cellular Biology, 13(12), 7913–7924. 10.1128/MCB.13.12.7913

Geiser, J. R., Van Tuinen, D., Brockerhoff, S. E., Neff, M. M., & Davis, T. N. (1991). Can calmodulin function without binding calcium? Cell, 65(6), 949–959. 10.1016/0092-8674(91)90547-C

Geli, M. I. (1998). Distinct functions of calmodulin are required for the uptake step of receptor-mediated endocytosis in yeast: The type I myosin Myo5p is one of the calmodulin targets. The EMBO Journal, 17(3), 635–647. 10.1093/emboj/17.3.635

Halling, D. B., Liebeskind, B. J., Hall, A. W., & Aldrich, R. W. (2016). Conserved properties of individual Ca^2+^ -binding sites in calmodulin. Proceedings of the National Academy of Sciences, 113(9). 10.1073/pnas.1600385113

Laplante, C., Berro, J., Karatekin, E., Hernandez-Leyva, A., Lee, R., & Pollard, T. D. (2015). Three Myosins Contribute Uniquely to the Assembly and Constriction of the Fission Yeast Cytokinetic Contractile Ring. Current Biology, 25(15), 1955–1965. 10.1016/j.cub.2015.06.018

Lee, W.-L., Bezanilla, M., & Pollard, T. D. (2000). Fission Yeast Myosin-I, Myo1p, Stimulates Actin Assembly by Arp2/3 Complex and Shares Functions with Wasp. Journal of Cell Biology, 151(4), 789–800. 10.1083/jcb.151.4.789

Lin, Y. M., Liu, Y. P., & Cheung, W. Y. (1974). Cyclic 3′: 5′-Nucleotide Phosphodiesterase. Journal of Biological Chemistry, 249(15), 4943–4954. 10.1016/S0021-9258(19)42412-5

Masuda, H., Mori, R., Yukawa, M., & Toda, T. (2013). Fission yeast MOZART1/Mzt1 is an essential γ-tubulin complex component required for complex recruitment to the microtubule organizing center, but not its assembly. Molecular Biology of the Cell, 24(18), 2894–2906. 10.1091/mbc.E13-05-0235

Mohanta, T. K., Kumar, P., & Bae, H. (2017). Genomics and evolutionary aspect of calcium signaling event in calmodulin and calmodulin-like proteins in plants. BMC Plant Biology, 17(1), 38. 10.1186/s12870-017-0989-3

Moreno, S., Klar, A., & Nurse, P. (1991). [56] Molecular genetic analysis of fission yeast Schizosaccharomyces pombe. In Methods in Enzymology (Vol. 194, pp. 795–823). Elsevier. 10.1016/0076-6879(91)94059-L

Moser, M. J., Flory, M. R., & Davis, T. N. (1997a). Calmodulin localizes to the spindle pole body of Schizosaccharomyces pombe and performs an essential function in chromosome segregation. Journal of Cell Science, 110(15), 1805–1812. 10.1242/jcs.110.15.1805

Moser, M. J., Flory, M. R., & Davis, T. N. (1997b). Calmodulin localizes to the spindle pole body of *Schizosaccharomyces pombe* and performs an essential function in chromosome segregation. Journal of Cell Science, 110(15), 1805–1812. 10.1242/jcs.110.15.1805

Moser, M. J., Lee, S. Y., Klevit, R. E., & Davis, T. N. (1995). Ca2+ Binding to Calmodulin and Its Role in Schizosaccharomyces pombe as Revealed by Mutagenesis and NMR Spectroscopy. Journal of Biological Chemistry, 270(35), 20643–20652. 10.1074/jbc.270.35.20643

Nyegaard, M., Overgaard, M. T., Søndergaard, M. T., Vranas, M., Behr, E. R., Hildebrandt, L. L., Lund, J., Hedley, P. L., Camm, A. J., Wettrell, G., Fosdal, I., Christiansen, M., & Børglum, A. D. (2012). Mutations in calmodulin cause ventricular tachycardia and sudden cardiac death. American Journal of Human Genetics, 91(4), 703–712. 10.1016/j.ajhg.2012.08.015

Perreault-Micale, C., Shushan, A. D., & Coluccio, L. M. (2000). Truncation of a Mammalian Myosin I Results in Loss of Ca2+-sensitive Motility. Journal of Biological Chemistry, 275(28), 21618–21623. 10.1074/jbc.M000363200

Poddar, A., Hsu, Y.-Y., Zhang, F., Shamma, A., Kreais, Z., Muller, C., Malla, M., Ray, A., Liu, A. P., & Chen, Q. (2022). Membrane stretching activates calcium permeability of a putative channel Pkd2 during fission yeast cytokinesis. Molecular Biology of the Cell, 33(14), ar134. 10.1091/mbc.E22-07-0248

Poddar, A., Sidibe, O., Ray, A., & Chen, Q. (2021). Calcium spikes accompany cleavage furrow ingression and cell separation during fission yeast cytokinesis. Molecular Biology of the Cell, 32(1), 15–27. 10.1091/mbc.E20-09-0609

Rhind, N., Chen, Z., Yassour, M., Thompson, D. A., Haas, B. J., Habib, N., Wapinski, I., Roy, S., Lin, M. F., Heiman, D. I., Young, S. K., Furuya, K., Guo, Y., Pidoux, A., Chen, H. M., Robbertse, B., Goldberg, J. M., Aoki, K., Bayne, E. H., … Nusbaum, C. (2011). Comparative Functional Genomics of the Fission Yeasts. Science, 332(6032), 930–936. 10.1126/science.1203357

Rosenberg, J. A., Tomlin, G. C., McDonald, W. H., Snydsman, B. E., Muller, E. G., Yates, J. R., & Gould, K. L. (2006). Ppc89 links multiple proteins, including the septation initiation network, to the core of the fission yeast spindle-pole body. Molecular Biology of the Cell, 17(9), 3793–3805. 10.1091/mbc.e06-01-0039

Rüthnick, D., Vitale, J., Neuner, A., & Schiebel, E. (2021). The N-terminus of Sfi1 and yeast centrin Cdc31 provide the assembly site for a new spindle pole body. Journal of Cell Biology, 220(3), e202004196. 10.1083/jcb.202004196

Stirling, D. A., Welch, K. A., & Stark, M. J. (1994). Interaction with calmodulin is required for the function of Spc110p, an essential component of the yeast spindle pole body. The EMBO Journal, 13(18), 4329–4342. 10.1002/j.1460-2075.1994.tb06753.x

Sundberg, H. A., Goetsch, L., Byers, B., & Davis, T. N. (1996). Role of Calmodulin and Spc110p Interaction in the Proper Assembly of Spindle Pole Body Components. The Journal of Cell Biology, 133.

Takeda, T., & Yamamoto, M. (1987). Analysis and in vivo disruption of the gene coding for calmodulin in Schizosaccharomyces pombe. Proceedings of the National Academy of Sciences, 84(11), 3580– 3584. 10.1073/pnas.84.11.3580

Tang, X., Huang, J., Padmanabhan, A., Bakka, K., Bao, Y., Yuelin Tan, B., Zacheus Cande, W., & Balasubramanian, M. K. (2011). Marker reconstitution mutagenesis: A simple and efficient reverse genetic approach. Yeast, 28(3), 205–212. 10.1002/yea.1831

Tomlin, G. C., Morrell, J. L., & Gould, K. L. (2002). The spindle pole body protein Cdc11p links Sid4p to the fission yeast septation initiation network. Molecular Biology of the Cell, 13(4), 1203–1214. 10.1091/mbc.01-09-0455

Win, T. Z., Gachet, Y., Mulvihill, D. P., May, K. M., & Hyams, J. S. (2001). Two type V myosins with non-overlapping functions in the fission yeast Schizosaccharomyces pombe: Myo52 is concerned with growth polarity and cytokinesis, Myo51 is a component of the cytokinetic actin ring. Journal of Cell Science, 114(1), 69–79. 10.1242/jcs.114.1.69

Wu, J.-Q., Kuhn, J. R., Kovar, D. R., & Pollard, T. D. (2003). Spatial and Temporal Pathway for Assembly and Constriction of the Contractile Ring in Fission Yeast Cytokinesis. Developmental Cell, 5(5), 723–734. 10.1016/S1534-5807(03)00324-1

